# Synaptopodin is required for efficient intercellular spread by bacterial pathogens

**DOI:** 10.1101/2023.04.25.537990

**Authors:** Jenna M. Vickery, Jody D. Toperzer, Julie E. Raab, Desmond J. Hamilton, Tucker B. Harju, Thao N. Huynh, Laurel L. Lenz, Liheng Zhou, Sean P. Colgan, Brian C. Russo

## Abstract

For many intracellular pathogens, their virulence depends on an ability to spread between cells of an epithelial layer. For intercellular spread to occur, these pathogens deform the plasma membrane into a protrusion structure that is engulfed by the neighboring cell. Although the polymerization of actin is essential for spread, how these pathogens manipulate the actin cytoskeleton in a manner that enables protrusion formation is still incompletely understood. Here, we show that butyrate responsive pathways promote intercellular spread by *Shigella flexneri*. We identify the mammalian actin binding protein synaptopodin, a butyrate responsive gene, as required for efficient intercellular spread of *S. flexneri* and *Listeria monocytogenes*. We show synaptopodin enhances the recruitment of actin to bacteria and stabilizes the actin tail. We show that, for *S. flexneri,* synaptopodin presence enables protrusions to form and to resolve at a greater rate, indicating that greater stability of the actin tail enables the bacteria to push against the membrane with greater force. We demonstrate that synaptopodin recruitment around bacteria requires the bacterial protein IcsA, and we show that this recruitment is further enhanced in a type 3 secretion system dependent manner. These data establish synaptopodin as required for intracellular bacteria to stabilize the actin cytoskeleton in a manner that enables efficient protrusion formation and identify for the first time that synaptopodin contributes to bacterial pathogenesis.

## Introduction

Many cytosolic-dwelling pathogens recruit actin-associated proteins to enable intercellular spread between mammalian cells (1), including *Listeria* spp*., Rickettsia* spp*., Burkholderia* spp*., Mycobacterium marinum, Shigella* spp., and poxviruses. The forces generated from the polymerization of actin enable their intracellular motility and movement between cells (2, 3). In order to polymerize actin, these organisms subvert preexisting mammalian cytoskeletal pathways. Whereas the ability to repurpose the cellular cytoskeleton is critical for their virulence, the molecular mechanisms that enable actin motility among these pathogens are still incompletely understood (4–6).

To gain new insights about how intracellular bacteria spread between cells, we focused on studying the model cytosolic pathogen *Shigella flexneri.* This gram-negative bacterium is a leading cause of bacillary dysentery, a form of bloody diarrhea (6). *S. flexneri* causes disease by invading cells of the colonic epithelium and by spreading between cells in this epithelial layer (7, 8). Cytosolic *S. flexneri* express IcsA at one bacterial pole (9); IcsA binds Neural Wiskott-Aldrich syndrome protein (N-WASP), which then recruits the Actin Related Protein 2/3 complex (Arp2/3) and promotes actin nucleation (10). The polymerization of actin enables the bacteria to move in the cytosol of the cell (11). At the periphery of the cell, some bacteria deform the plasma membrane into a protrusion structure that pushes into the neighboring (recipient) cell (12), where it is actively engulfed. In the recipient cell, these protrusions are resolved into double membrane vacuoles. The bacteria quickly escape the double membrane vacuole into the cytosol, where they can repeat this cycle of intercellular spread (13). Intercellular spread allows the bacterium to access nutrients while evading the immune system, and both the invasion of the colonic epithelium and the spread between cells are essential for pathogenesis (5, 14–17). Defining the molecular mechanisms that promote the intercellular spread of *Shigella* is essential to understand how this organism causes disease.

Butyrate is produced by the microbiota and is rapidly metabolized by intestinal epithelial cells (18–20). In these cells, butyrate results in a variety of changes, as it is a potent histone deacetylase (20–22). The pathways regulated by butyrate include a variety of cellular processes, such as energy consumption, hypoxia inducible factors, wound healing, inflammation, and barrier function. We reasoned that many of the targets of butyrate could be relevant during infection and that we could leverage the cellular response to butyrate as a way to uncover new genes and pathways relevant for infection.

Here, we show that butyrate promotes *S. flexneri* intercellular spread in intestinal epithelial cells. Based on previous RNAseq studies, we show that synaptopodin, a gene upregulated by butyrate, is important for intercellular spread of *S. flexneri* and *L. monocytogenes.* We show protrusion formation and actin tail stability depend on synaptopodin presence. Synaptopodin is recruited to bacteria in a manner that is dependent on the bacterial protein IcsA, and its recruitment is additionally enhanced by a Spa15-dependent bacterial protein. We show that synaptopodin function likely occurs in a manner distinct from previously described pathways. Together, these data, using *S. flexneri* as a model, identify synaptopodin as a critical cellular component that enables efficient intercellular spread by cytosolic pathogens.

## Results

### Synaptopodin is required for efficient intercellular spread by *S. flexneri*

To identify new proteins and pathways required for intercellular spread, we manipulated cellular function by adding the short chain fatty acid, butyrate, to the culture media. We compared the effect of butyrate on the efficiency of spread for *S. flexneri* through a monolayer of human colonic adenocarcinoma epithelial (T84) cells using a plaque assay. In this assay, infected cells are killed by the bacteria as they spread, creating clearings in the monolayer, known as “plaques”. The number of plaques is a function of the efficiency of invasion, and the area of the plaque is a function of the efficiency of intercellular spread. We observed larger plaques in the presence of butyrate (Fig. 1A-B), demonstrating butyrate enables more efficient spread by *S. flexneri* in these cells. On its own, butyrate did not change the replication of *S. flexneri* (Fig. S1), suggesting that the increased efficiency of spread is independent of bacterial replication and that butyrate may alter cellular pathways necessary to promote intercellular spread.

**Figure 1:**
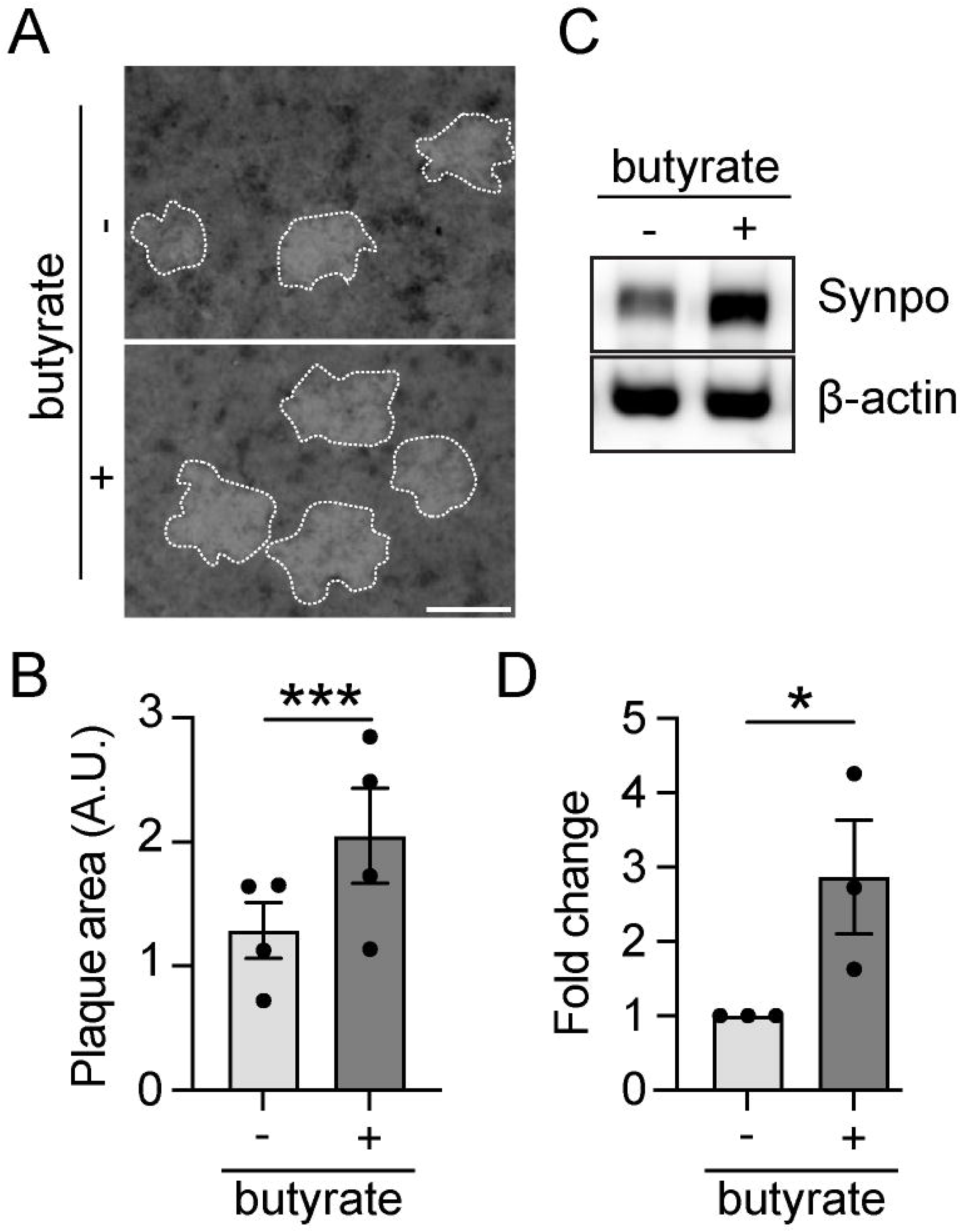
Butyrate promotes efficient *S. flexneri* intercellular spread. (A-B) Monolayers of T84 cells were infected with *S. flexneri* in the presence of 0 or 5 mM butyrate, stained with neutral red at 48 hours of infection, and imaged. (A) Representative images. Dotted lines show plaque boundary; scale bar, 0.6 mm. (B) Quantification of the area of plaques from images represented in A, >50 plaques were measured per condition per experiment. Dots are independent experiments. (C,D) T84 cells were treated with 0 or 5 mM butyrate for 24 hours, and Synaptopodin abundance was determined with western blot. (B,D) ***, p<0.001; *, p<0.05 by paired t-test. Data are mean ± SEM.

Previous RNA sequencing of T84 cells identified genes whose expression was altered by butyrate and identified Synaptopodin (23). We show that butyrate caused a 3-fold increase in synaptopodin protein in our assays (Fig. 1C-D). Synaptopodin is an actin binding protein that alters the efficiency of actin polymerization and interacts with cell-cell junction proteins. Its robust induction by butyrate, its subcellular localization at sites of intercellular spread, and its regulation of actin-dependent processes indicated that synaptopodin could have roles during *S. flexneri* intercellular spread.

To investigate this, a control or synaptopodin specific shRNA was introduced into T84 cells, and the efficiency of intercellular spread was determined by a plaque assay. Synaptopodin protein abundance was reduced in the presence of the synaptopodin-specific shRNA (23), and *S. flexneri* formed smaller plaques in Synpo knockdown cells compared to control cells (Fig. 2A-B). These data suggest that synaptopodin promotes intercellular spread by *S. flexneri*.

**Figure 2:**
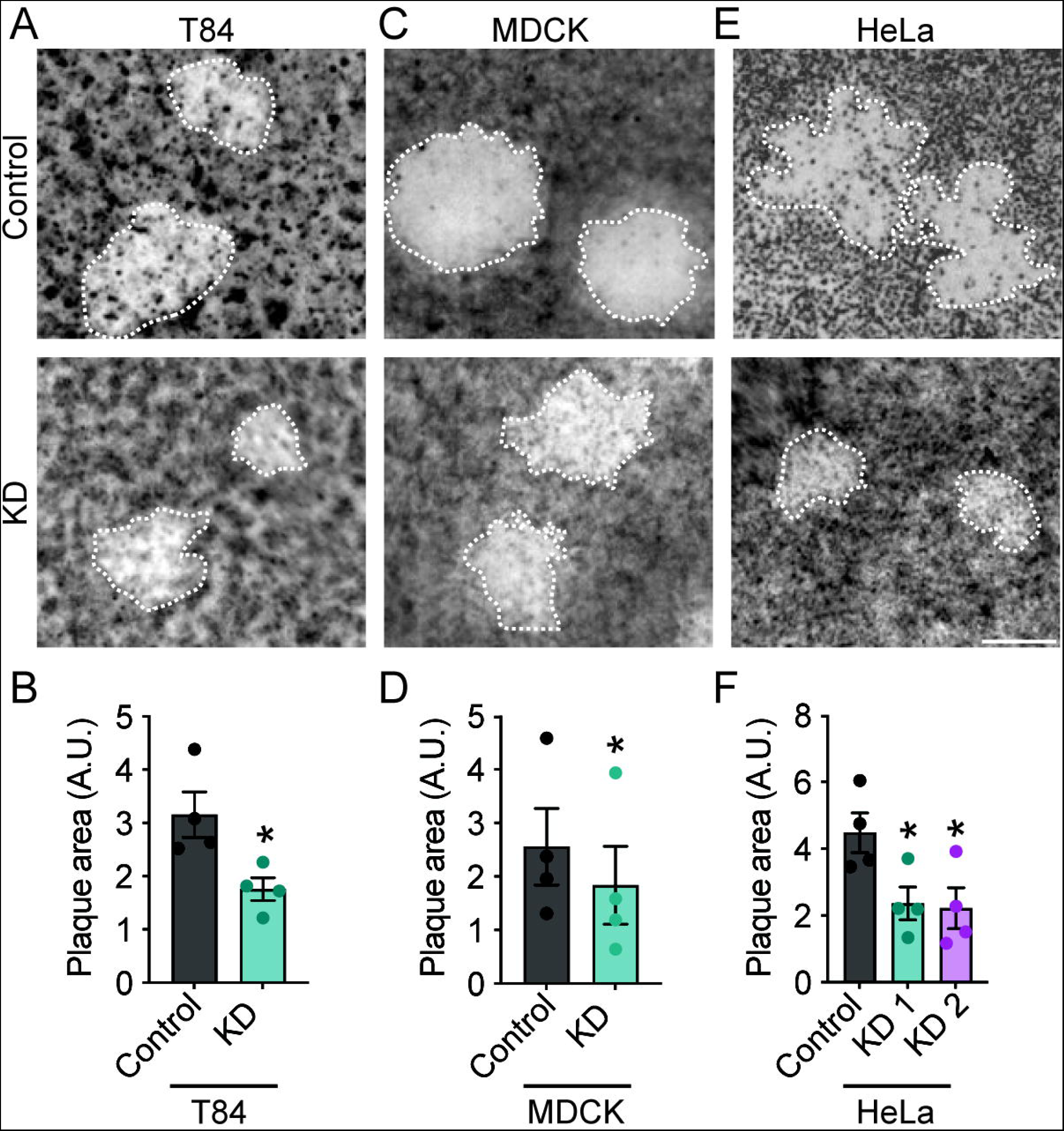
Synaptopodin is required for efficient intercellular spread of *S. flexneri*. Monolayers of T84 (A-B), MDCK (C-D), or HeLa (E-F) cells that produce a control or synaptopodin specific shRNA were infected with *S. flexneri* and stained with neutral red at 48 hours of infection. (A, C, and E) Representative images. Dotted lines indicate plaque boundary. Scale bar, 0.5 mm. (B, D, and F) Quantification of the area of plaques from images represented in A, C, and E. 15-60 plaques measured per condition per experiment. Dots are independent experiments. *, p<0.05 by paired t-test (B and D) or by one-way ANOVA with Dunnett’s *post hoc* test (F). Data are mean ± SEM.

The effect of synaptopodin on intercellular spread was not specific to T84 cells; plaques were significantly smaller in Madin-Darby canine kidney (MDCK) cells producing a control shRNA compared to MDCK cells producing a synaptopodin-specific shRNA (Fig. 2C-D). Synaptopodin was efficiently knocked-down in these cells [Fig. S2A-B, (23, 24)]. The ability of *S. flexneri* to spread through monolayers of HeLa cells producing a control shRNA or one of five synaptopodin-specific shRNAs was also investigated. Plaques formed in monolayers of HeLa cells producing four of the five synaptopodin-specific shRNAs were smaller than plaques formed in cells producing the control shRNA (Fig. 2E-F and S2C-D). The four cell lines displaying reduced plaque size also demonstrated a reduction in synaptopodin protein level (Fig. S2E-F). Thus, the smaller plaque size observed above was unlikely the result of non-specific effects of an individual shRNA. *S. flexneri* replicated at a similar efficiency in T84, MDCK, and HeLa cells that produced or lacked synaptopodin (Fig. S2G-L), indicating that the impact of synaptopodin on plaque size is not due to differences in bacterial replication. Moreover, since similar results were observed in T84, MDCK, and HeLa cells, these data indicate that the requirement of synaptopodin for *S. flexneri* spread is unlikely to be specific for a particular cell type.

*L. monocytogenes* is a gram-positive bacterium that, like *S. flexneri,* requires actin-based motility for intercellular spread (25). The ability of *L. monocytogenes* to spread through a monolayer of HeLa cells producing a control shRNA was compared to its ability to spread through cells producing a synaptopodin-specific shRNA by fluorescence microscopy. Similar to the observations with *S. flexneri,* plaques formed by *L. monocytogenes* were larger in the presence of synaptopodin (Fig. S3), demonstrating that synaptopodin is required for intercellular spread of *Listeria*. Together, these data show that *S. flexneri* and *L. monocytogenes* require synaptopodin for efficient intercellular spread and indicate the requirement for synaptopodin may be a general feature of intracellular bacterial pathogens that spread.

### Synaptopodin is required for protrusion formation by *S. flexneri*

To investigate how synaptopodin contributes to efficient intercellular spread, we focused on aspects of the bacterial intracellular life cycle that require actin. We assessed the ability of bacteria to form plasma membrane protrusions in HeLa cells producing either a control or a synaptopodin-specific shRNA. We determined the percentage of bacteria present in a protrusion at 90 minutes of infection by examining the manner of actin polymerized at the posterior pole of bacteria by immunofluorescence microscopy. We performed similar studies in which a membrane-anchored YFP was introduced into cells and used to mark the plasma membrane. For both approaches, a smaller percentage of bacteria in synaptopodin knockdown cells were present in plasma membrane protrusions compared to the percentage of bacteria in protrusions in control cells (Fig. 3A-B and S4), demonstrating that synaptopodin is required for efficient protrusion formation.

**Figure 3:**
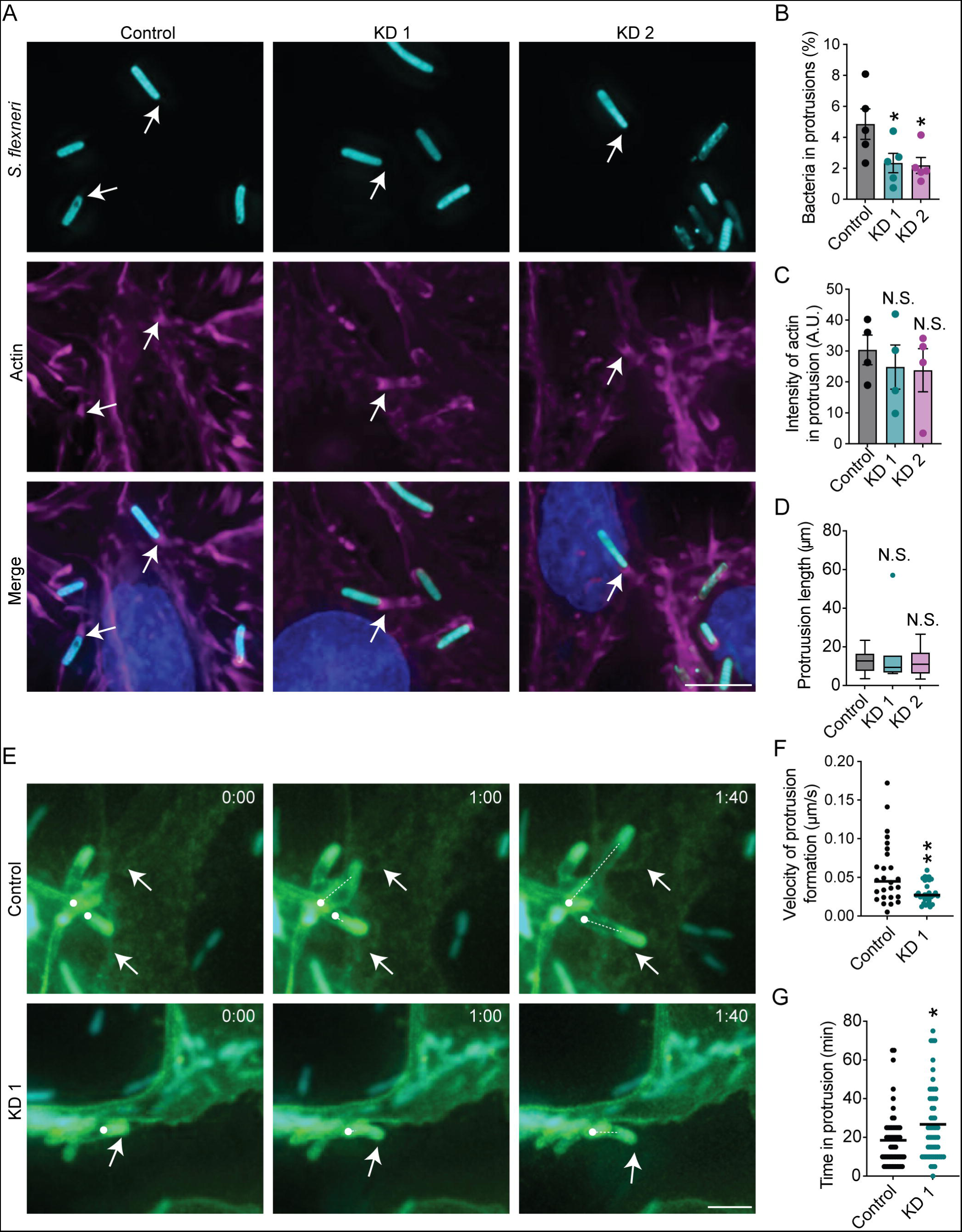
Synaptopodin is required for protrusion formation by *S. flexneri*. (A) Representative images of HeLa cells infected with *S. flexneri.* Cyan, *S. flexneri;* magenta, actin; blue, DNA. Arrows, bacteria in protrusions. Scale bar, 5 µM. 5-10 fields were analyzed per condition per experiment. (B) Quantification of the percentage of intracellular bacteria within a protrusion. (C) Quantification of the intensity of actin in protrusions. (B and C) Dots are independent experiments. (D) Quantification of the length of protrusions. Tukey method used to define whiskers. (B-D) N.S., not significant; *, p<0.05 by one-way ANOVA with Holm-Sidak’s *post hoc* test. (E-G) Time lapse images from videos of bacteria forming plasma membrane protrusions. (E) Representative images of protrusion formation. Cyan, S. *flexneri;* green, plasma membrane. Scale bar 5 µM. Arrows, bacteria in protrusions; dot, protrusion origination point; dotted line, progression of protrusion formation. (F) Quantification of the speed of protrusion formation from videos represented in E. (G) Quantification of the duration bacteria remain in protrusions. (F-G) Dots are individual bacteria, pooled from 3 independent experiments. *, p<0.05 by Student’s t-test. Data are mean ± SEM.

Synaptopodin is associated with the bundling and elongation of actin filaments in cells (26, 27), but actin intensity at the plasma membrane near protrusions was similar in the presence and absence of synaptopodin (Fig. 3C), indicating that the protrusion defect is not due to defects in the amount of actin recruited to protrusions. Protrusion formation is also dependent on actin that is polymerized by formins. The formin mDia1 is required for formation of protrusions by *S. flexneri* and *L. monocytogenes* (28, 29). mDia1 was efficiently recruited to protrusions formed in control or synaptopodin knockdown cells (Fig. S5), indicating reduced mDia1 levels around bacteria are not driving the reduction in protrusion formation in the relative absence of synaptopodin. The length of the protrusion is more generally associated with the efficiency of intercellular spread (30). Protrusion lengths were similar irrespective of synaptopodin levels in HeLa cells (Fig. 3D), indicating that the requirement for synaptopodin was not associated with protrusion length.

We next investigated the kinetics of protrusion formation by *S. flexneri* to further clarify the role of synaptopodin in the development of membrane protrusions. To visualize the formation of protrusions, intracellular bacteria were tracked by time-lapsed microscopy during infection of HeLa cells that produce a YFP that is anchored to the plasma membrane (31). Bacteria in control cells formed protrusions at a greater velocity than bacteria in cells lacking synaptopodin (Fig. 3E-F). Protrusions formed in the presence of synaptopodin were also faster to resolve, lasting an average of 18 min in control cells compared to 27 min in synaptopodin knockdown cells (Fig. 3G). These data show synaptopodin enables more rapid formation and resolution of protrusions.

### Synaptopodin regulates actin tail formation by *S. flexneri*

The ability to efficiently assemble an actin tail in the cell cytosol is associated with efficient protrusion formation at the plasma membrane (32). We hypothesized that the defects in protrusion formation were the result of aberrant actin tail formation. To test this, we infected HeLa cells with *S. flexneri* and characterized the actin tails. Although a similar percentage of bacteria formed actin tails in the control and synaptopodin knockdown cells (Fig. 4A-B), actin tails that formed in the synaptopodin knockdown cells were significantly shorter than actin tails in the control cells (Fig. 4C). In MDCK cells, the actin tails formed by *S. flexneri* in the control cells were also significantly longer than tails in the synaptopodin knockdown cells (Fig. S6A-B). Previous studies have shown that the length of the actin tails correlates with the rate of actin polymerization and the speed bacteria move in the cytosol (33). Since we observed shorter tails in HeLa and MDCK cells, it suggested to us that the rate of actin polymerization could be reduced in the absence of synaptopodin and would result in *S. flexneri* moving more slowly through the cytosol of cells. To test this, time-lapsed microscopy was performed on intracellular bacteria in HeLa cells to evaluate the efficiency of actin-based motility. In contrast to our expectation based on the length of the actin tails, the bacterial velocity was similar in the control and synaptopodin knockdown cells (Fig. 4D-E). Since bacteria were moving at a similar speed, it suggested that the rate of actin polymerization was similar in the presence and absence of synaptopodin and indicated that synaptopodin stabilizes the polymerized actin tail resulting in longer actin tails in the control cells.

**Figure 4:**
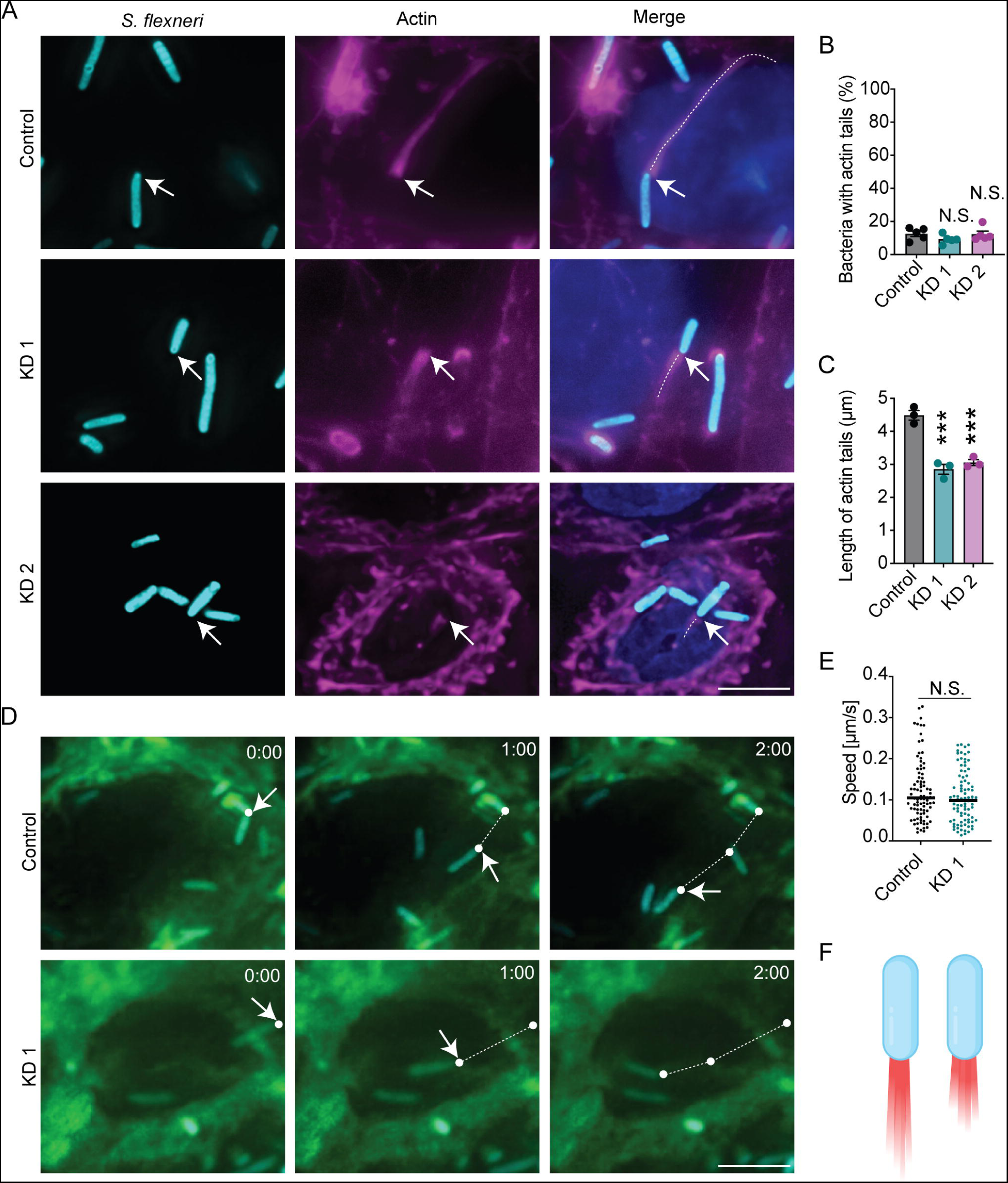
Synaptopodin regulates actin tail formation by *S. flexneri*. Infection of HeLa cells infected with *S. flexneri* for 90 min. (A) Representative images. Cyan, *S. flexneri;* magenta, actin; blue, DNA. Arrows, bacteria with actin tails. Dotted line, trace of actin tail. Scale bar, 5 µM. (B) Quantification of percentage of bacteria with actin tails. (C) Quantification of the length of actin tails. (D-E) Time-lapsed microscopy of HeLa cells producing a membrane anchored YFP infected with *S. flexneri* producing CFP. (D) Representative still images from time-lapsed videos for 2 minutes from initial observation of motility. Cyan, *S. flexneri*; green, plasma membrane. Dots are the position of bacteria at each time point, dotted line is arbitrary straight line between dots showing bacterial progression over time. Scale bar, 5 µM. (E) Quantification of the speed at which bacteria move in the cytosol from videos represented in E. Mean indicated by bar. (B, C) 5-10 fields were analyzed per experiment. Dots are independent experiments. Data are mean ± SEM. N.S., not significant; *, p<0.05; **, p<0.1; ***, p<0.001 by one-way ANOVA with Dunnett’s *post hoc* test. (E) N.S., not significant; *, p<0.05 by Student’s t-test. Dots are individual bacteria pooled together from three experiments. (F) Schematic depicting shorter tails are formed in cells lacking synaptopodin compared to tails formed in control cells. Generated with BioRender.

In addition to differing in length, the width of the actin tails at the base of the bacteria was wider in control HeLa cells than in synaptopodin knockdown HeLa cells (Fig. S7A-B). Interestingly, actin tails at the base of the bacteria in MDCK cells were not significantly different in width between control and synaptopodin knockdown cells (Fig. S6C). We also noted additional phenotypic changes to the tails in HeLa cells lacking synaptopodin. Polymerized actin was more frequently present along the lateral sides of the posterior end of bacteria in the control cells compared to in synaptopodin knockdown cells (Fig. S7C). During bacterial infection or when examining actin polymerization around lipid vesicles, the actin on the sides of the bacteria or vesicle creates an inward force at the base that is predicted to contribute to the propulsive force (34, 35). Seeing that there was no difference in the velocity at which *S. flexneri* traveled whether or not actin extended along the sides of the bacteria, we hypothesized that the additional actin may stabilize the tail and keep the actin tail in line with the bacteria, which we anticipated would stabilize the path along which the bacteria travels. To investigate this, we measured the angle at which the actin tail extended from the posterior pole of the bacteria. We set actin tails that formed completely in line with the longitudinal axis of the bacteria at 0° and deviations from this alignment resulted in an increased angle. In the synaptopodin knockdown cells, the actin tails were less aligned with the bacteria’s longitudinal axis than the tails associated with bacteria in the control cells, which resulted in a greater angle from the longitudinal axis for bacteria in synaptopodin knockdown cells (Fig. S7D). These data show that synaptopodin enables actin presence along the lateral sides near the posterior end of the bacteria and that synaptopodin enables the actin tail to be more in line with the bacteria. We theorized that the angle of the actin tail would decrease the forward momentum that the actin nucleation creates at the base of the bacteria. We used the formula, [*sin (angle of the tail(θ)) + cos (angle of the tail(θ))*], to calculate the total force of the actin tail on the bacteria. We then calculated the percentage of force that contributed to the forward momentum of the bacteria by the formula, [*(cos (angle of the tail(θ)) / total force) * 100*]. The percentage of the theoretical forward movement of bacteria in the control cells is higher than in the synaptopodin knockdown cell lines (Fig. S7E). We hypothesized that the defects in the actin tail length, width, organization, and angle of attachment would lead to a defect in the linear trajectory of the bacteria in the absence of synaptopodin. We tracked intracellular bacteria with time-lapsed microscopy to evaluate the linear trajectory. We calculated the distance an individual bacteria traveled and divided it by the distance of the theoretical most efficient path – a straight line. A bacterium that took the most efficient path would have a linear trajectory ratio of 1.0. We observed that bacteria deviated further from an ideal path in cells lacking synaptopodin compared to control cells (Fig. S7F), which showed that bacteria traveled along a straighter path in the presence of synaptopodin.

We asked whether the defects associated with the actin tails were specific to *S. flexneri* or a more general result of pathogens that spread between cells with lower levels of synaptopodin. We tested whether the length of actin tails was similarly decreased during *L. monocytogenes* infection of HeLa cells. Similar to *S. flexneri,* actin tails formed by *L. monocytogenes* were longer in the presence of synaptopodin compared to the actin tails formed in the absence of synaptopodin (Fig. S8).

Altogether, these data show that synaptopodin presence stabilizes polymerized actin tails resulting in longer actin tails and that, in HeLa cells, synaptopodin additionally enables actin accumulation at the bacterial pole in a manner that generates more directed forward momentum.

### Synaptopodin colocalizes around *S. flexneri*

Based on the requirement for synaptopodin in organizing the actin tail and in forming protrusions, we hypothesized that *S. flexneri* would recruit synaptopodin to its pole during intracellular infection. To test this, we infected HeLa and MDCK cells with *S. flexneri* and used immunofluorescence microscopy to visualize synaptopodin localization in cells. Synaptopodin was recruited to bacteria in the cytosol (Fig. S9A-C and S10). This recruitment of synaptopodin mirrored that of actin and was observed at the poles, the lateral sides of the bacteria, or both. Furthermore, synaptopodin localizes to bacteria that also displayed actin recruitment, but synaptopodin was not observed around bacteria that failed to recruit actin (Fig. S9-S10). These data show that synaptopodin is recruited to bacteria that nucleate actin polymerization.

Knockdown of synaptopodin caused fewer bacteria to recruit synaptopodin (Fig. S9A-B), and among the bacteria that showed synaptopodin recruitment, there was a reduction in the intensity of synaptopodin (Fig. S9C). Although loss of synaptopodin did not reduce the percentage of bacteria that formed an actin tail (Fig. 4B), the knockdown of synaptopodin significantly reduced the number of bacteria that recruited actin (Fig. S9D), yet there was not a significant difference in the intensity of the actin signal among the bacteria that recruited actin in the presence or absence of synaptopodin in these experiments (Fig. S9E). These data show that synaptopodin is recruited to the poles and sides of intracellular bacteria that polymerize actin and synaptopodin presence enhances the number of bacteria that recruit actin.

### Synaptopodin promotes myosin IIb recruitment around *S. flexneri*

The localization of IcsA to the posterior pole of *S. flexneri* results in the recruitment of N-WASP and the Arp2/3 complex and enables the initiation of actin polymerization and bacterial motility (10). Synaptopodin is known to regulate this process through another protein Nck1 (36). Synaptopodin promotes Nck1 abundance in cells by blocking c-Cbl-mediated ubiquitination of Nck1, and Nck1 assists in N-WASP and Arp2/3 recruitment during vaccina virus infection (36, 37). We tested whether the defect in actin recruitment in cells lacking synaptopodin was due to a disruption in this synaptopodin-Nck1-Arp2 pathway. We observed a similar percentage of bacteria that recruited Arp2, and these bacteria were associated with similar levels of Arp2 as measured by the intensity of the fluorescent signal of Arp2 around bacteria (Fig. S11A-C). We tested whether Nck1 was recruited to the bacteria during intracellular infection. We found no evidence of Nck1 recruitment in our cells (Fig. S11D). We examined the ability of *S. flexneri* to recruit N-WASP and observed a similar percentage of bacteria recruited N-WASP (Fig. S11E-F). Among the bacteria that recruited N-WASP, the amount of N-WASP was similar irrespective of synaptopodin levels (Fig. S11G). Consistent with our findings above that Synaptopodin likely stabilizes the polymerized actin in the tail rather than altering the rate of actin polymerization, here, we did not see defects in the recruitment of key proteins required to intiate actin polymerization at the posterior pole of the bacteria.

We further sought to examine whether we could identify interactions that were important for the formation of the protrusion. Synaptopodin is described to interact with non-muscle myosin IIb (myosin IIb) and α-actinin-4 at the plasma membrane, where together they form the contractomere complex that generates force so as to enable the extrusion of cells from a monolayer (24). Recent work shows that α-actinin-4 is present at actin structures formed during *S. flexneri* infection (1). We tested for the recruitment of myosin IIb and α-actinin-4 by immunofluorescence microscopy, and observed both were efficiently recruited to *S. flexneri.* Interestingly, a smaller percentage of bacteria recruited myosin IIb in the absence of synaptopodin (Fig. 5A-B), but among the bacteria that recruited myosin IIb, there was a similar abundance of myosin IIb as measured by fluorescent intensity irrespective of synaptopodin levels (Fig. 5C). We observed similar levels of α-actinin-4 recruited to bacteria and similar abundance of α-actinin-4 around the bacteria as measured by fluorescent intensity in control and synaptopodin knockdown cells (Fig. 5D-F). Altogether, these data indicate that synaptopodin presence enhances myosin IIb recruitment around *S. flexneri*.

**Figure 5:**
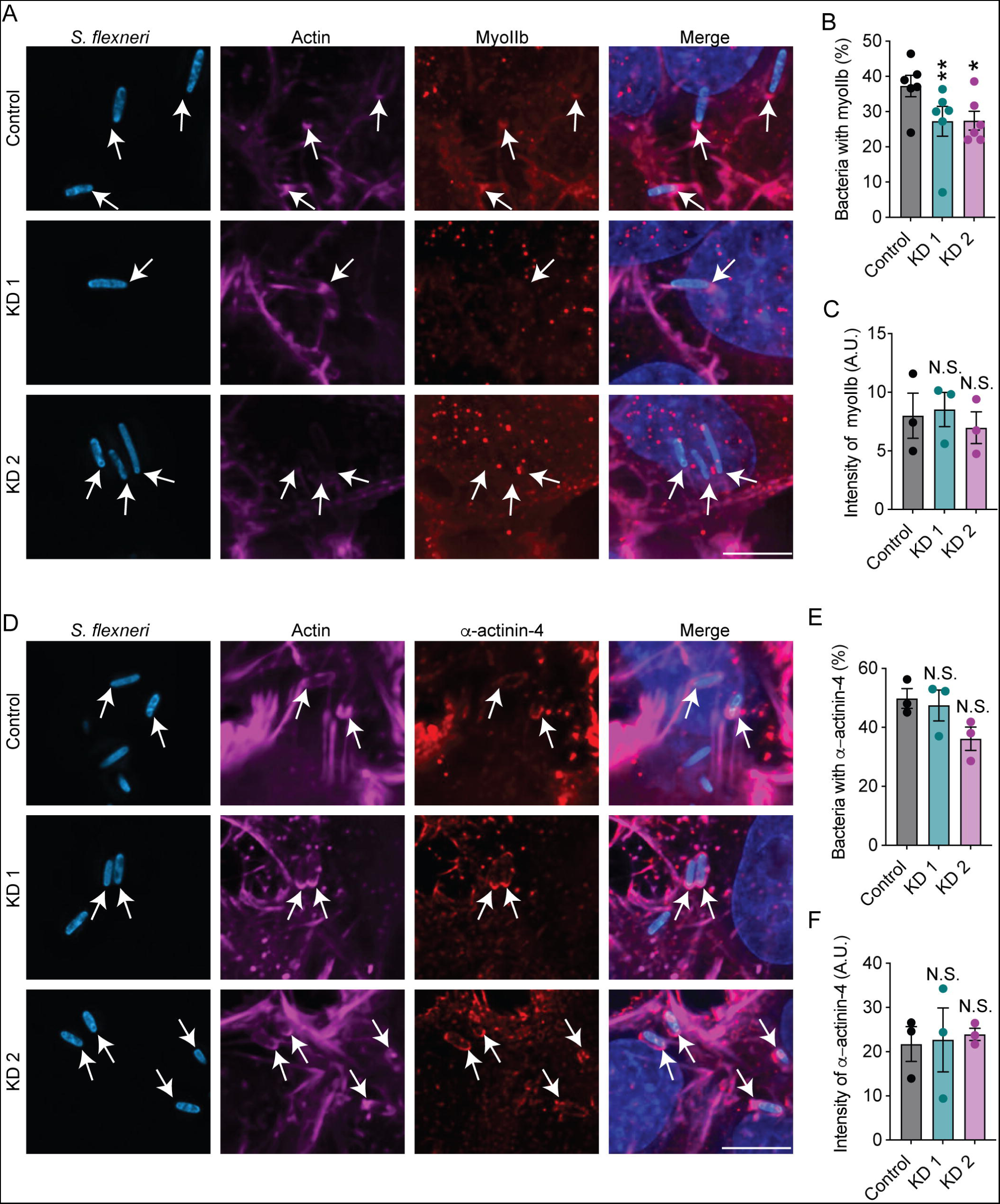
Synaptopodin is required for myosin IIb recruitment around bacteria. HeLa cells were infected with *S. flexneri* for 90 min. (A) Representative images. Cyan, *S. flexneri*; magenta, actin; red, myosin IIb; blue, DNA. Arrows, bacteria with myosin IIb. Scale bar, 5 µM. (B) Quantification of percentage of bacteria with myosin IIb. (C) Quantification of the intensity of myosin IIb around bacteria. (D) Representative images depicting α-actinin-4 recruitment around bacteria. Cyan, *S. flexneri;* magenta, actin; red, α-actinin-4; blue, DNA. Arrows, bacteria with α-actinin-4. Scale bar, 5 µM. (E) Quantification of the percentage of bacteria with α-actinin-4. (F) Quantification of the intensity of α-actinin-4. (B-C and E-F) 5-10 fields were analyzed per condition per experiment. Dots are independent experiments. Data are mean ± SEM. NS, not significant; *, p<0.05; **, p<0.01 by one-way ANOVA with Dunnett’s *post hoc* test.

### IcsA and the type 3 secretion system regulate synaptopodin recruitment around *S. flexneri*

*S. flexneri* requires a type three secretion system (T3SS) for intercellular spread (38–40). This type 3 secretion system delivers over 30 bacterial virulence proteins (effectors) into the cell (41). The delivered effectors coopt cellular signaling pathways and enable *S. flexneri* to thrive in the intracellular environment. Here, we sought to determine whether synaptopodin was recruited around the bacteria indirectly due to the high concentrations of polymerized actin around the bacteria, as a result of direct recruitment by *S. flexneri* secreted proteins, or both. To do so, we tested whether IcsA or the type 3 secreted effector proteins contributed to synaptopodin recruitment by investigating synaptopodin localization during cellular infection by several bacterial mutants: *S. flexneri* △*icsA* is unable to polymerize actin; *S. flexneri* △*spa15* lacks a chaperone for nine type 3 secreted effector proteins – the absence of Spa15 results in these proteins being unstable and degraded (42); *S. flexneri* △*mxiE* lacks the MxiE transcription factor that is necessary to transcriptionally activate the expression of more than 25 effectors – in the absence of MxiE these effectors are not produced in the bacteria (43–45). Together, the absence of Spa15 and MxiE enabled us to examine whether more than 30 effectors contribute to synaptopodin recruitment as two pooled groups. We infected HeLa cells with these mutants and quantified the percentage of bacteria that showed recruitment of synaptopodin and/or actin, and we measured the intensities of these recruited proteins. As expected, *S. flexneri* △*icsA* did not recruit actin or form actin tails during intracellular infection (Fig. 6A-B). Interestingly, bacteria lacking IcsA also did not recruit synaptopodin (Fig. 6C). A similar percentage of wild-type bacteria and bacteria lacking MxiE recruited actin and synaptopodin (Fig. 6A-C). Loss of Spa15 resulted in a smaller percentage of bacteria that recruited actin compared to wild-type bacteria, but a similar percentage of wild-type and Spa15 deficient bacteria recruited synaptopodin (Fig. 6C). When we examined the amount of actin and synaptopodin recruited around bacteria by comparing the intensity of fluorescent signal, we observed that except for the *icsA* mutant, actin was efficiently recruited around bacteria (Fig. 6D). Yet, when we examined the amount of synaptopodin around bacteria, we observed that bacteria lacking Spa15 or IcsA recruited less synaptopodin than wildtype bacteria (Fig. 6E). Altogether these data show that bacterial recruitment of synaptopodin is dependent on *S. flexneri* protein IcsA and is enhanced in a type 3 secretion-dependent manner.

**Figure 6:**
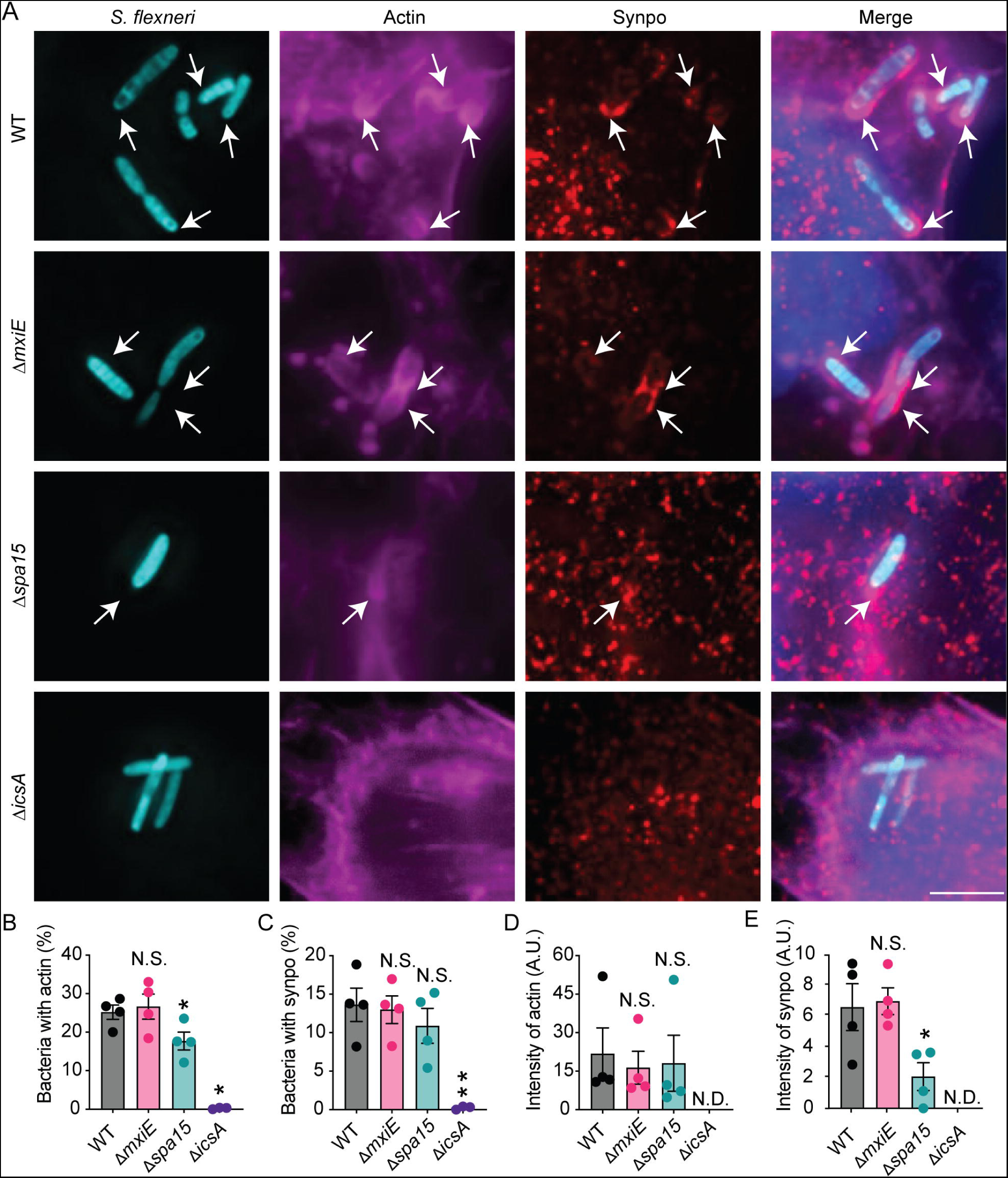
Synaptopodin is recruited around *S. flexneri* in an IcsA and a type 3 secretion system dependent manner. Control HeLa cells were infected for 90 min with *S. flexneri* strains. (A) Representative images. Cyan, *S. flexneri*; magenta, actin; red, synaptopodin (synpo); blue, DNA. Arrows, bacteria with actin and synaptopodin. Scale bar, 5 µM. (B) Quantification of the percentage of bacteria with actin. (C) Quantification of the percentage of bacteria with synaptopodin. (D) Quantification of the intensity of actin around bacteria. (E) Quantification of intensity of synaptopodin around bacteria. (B-E) 5-10 fields were analyzed per condition per experiment. Dots are independent experiments. Data are mean ± SEM. N.D., not detected. NS, not significant; *, p<0.05; **, p<0.01 by one-way ANOVA with Dunnett’s *post hoc* test.

## Discussion

Many cytosolic-dwelling pathogens require the subversion of actin and host proteins for intracellular motility. These pathogens remodel the cytoskeleton and plasma membrane resulting in the formation of protrusions, which facilitate intercellular spread. Here, we show that butyrate responsive pathways enhance *S. flexneri* intercellular spread. We identify that the butyrate responsive protein synaptopodin promotes *S. flexneri* and *L. monocytogenes* intercellular spread. We show that synaptopodin function is required to stabilize actin tails and to promote efficient protrusion kintecs, and we show that synaptopodin is recruited to bacteria in a manner that is dependent on the bacterial protein IcsA and the type 3 secretion system. Thus, these findings identify that bacterial pathogens, such as *S. flexneri* and *L. monocytogenes,* drive changes in the subcellular localization of synaptopodin in a manner that promotes actin tail stability, protrusion formation, and intercellular spread (Fig. 7).

**Figure 7.**
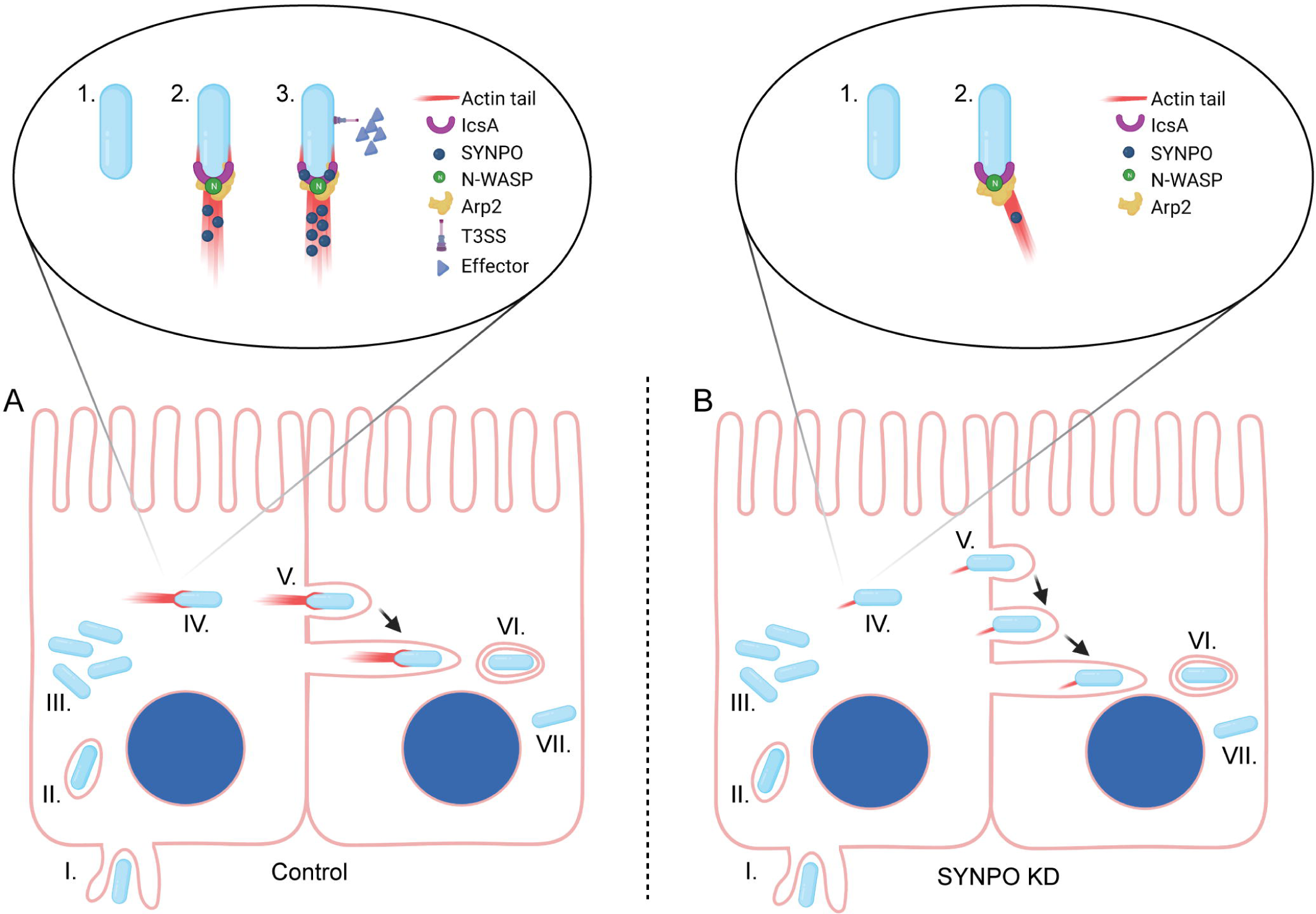
Schematic of *S. flexneri* intercellular spread in control versus synaptopodin knockdown cell lines. Schematic of *S. flexneri* intercellular lifecycle. (I) *S. flexneri* induces its uptake into cells. (II) Following uptake, it is present in a vacuole. (III) It escapes the vacuole into the cytosol where the bacteria replicate. A portion of the bacteria nucleate the polymerization of actin, which enables the bacteria to move in the cell cytosol. (V) Bacteria near the cell periphery deform the plasma membrane into a protrusion. (VI) The protrusion resolves into a double membrane vacuole. (VII) The bacteria escape the double membrane vacuole into the cytosol. (A) Cells with synaptopodin enabled bacteria to form actin tails that are longer and more aligned with the bacteria than cells lacking synaptopodin. Bacteria in cells with synaptopodin form protrusions at a higher velocity that resolve more quickly (V). (B) In cells lacking synaptopodin, the actin tails are shorter. Protrusions are formed at a slower velocity and take longer to resolve into a double membrane vacuole (V). (A-B) 1. Bacteria lacking IcsA do not polymerize actin or recruit synaptopodin. 2. Polar localization of IcsA is associated with synaptopodin recruitment, and IcsA recruits N-WASP and Apr2/3 irrespective of synaptopodin abundance in cells. 3. Synaptopodin recruitment to bacteria is further enhanced in a type 3 secretion dependent manner. Created with biorender.com

Here, we used butyrate as a tool to manipulate epithelial cell function. Whereas it is unclear whether this short chain fatty acid would have direct effects on intercellular spread during intestinal infection, there is reason to belive that *S. flexneri* infection would create conditions where butyrate responsive pathways in epithelial cells are relevant for spread. Butryate treatment is known to stabilize hypoxia indicble factor 1a (HIF1a) (22, 46). HIF is degraded under normoxia, but is stabilized and active under hypoxic conditions. *S. flexneri* infection is known to create hypoxic conditions in the intestinal epithelium (47, 48). Moreover, the stabilization of HIF is associated with increased levels of synaptopodin. We speculate that a consequence of *S. flexneri* induced hypoxia is the stabilization of HIF and upregulation of synaptopodin such that intercellular spread becomes more efficient.

Previous studies have shown that the length of the actin tail formed by *L. monocytogenes* directly correlates with the rate of actin polymerization and the intracellular velocity of the bacteria (33). Although synaptopodin presence enabled longer actin tails to form (Fig. 4C, S6, and S8), synaptopodin loss did not alter the speed the bacteria moved in the cytosol (Fig. 4D-E). Moreover, proteins associated with actin nucleation, such as N-WASP and Arp2, were efficiently recruited to bacteria irrespective of synaptopodin (Fig. S11A-C and E-G). These findings suggest that rather than altering the rate of actin polymerization, synaptopodin stabilizes the actin filaments that are already formed leading to the generation of longer actin tails. Similar to our findings, the stabilization of F-actin by synaptopodin enables the formation of dendritic spines in neurons (27, 49). Although the bacterial velocity is associated with actin monomer addition in the cytosol, it is likely that the rate of addition of new actin monomers as well as the rate of loss of old actin monomers may both contribute to the speed at which the protrusions form. Consistent with our model, the tail length did not correlate with bacterial speed in the cytosol, but was asocciated with a decreased rate of protrusion formation in the absence of synaptopodin (Fig. 3F).

Along with shorter actin tails, bacterial tails formed in the synaptopodin knockdown HeLa cells were less aligned with the longitudinal axis of the bacteria and lacked actin recruitment along the sides of the bacteria. Here, our data indicate that the additional actin provides a stabilizing effect by maintaining actin polymerization in line with the bacterial longitudinal axis and enabling the bacteria to move in a more directed manner forward (Fig. S7F). In MDCK cells, synaptopodin was associated with decreased length of actin tails, but not the width (Fig. S6). Our data indicate that a reduction of actin on the sides of the bacteria does not affect the velocity of cytosolic bacteria (Fig. 4). Rather, we speculate that this specific synaptopodin-dependent localization of actin in HeLa cells may be necessary at the membrane where the bacteria experience additional forces during the formation of the protrusion.

Previous studies have shown that synaptopodin forms a force-generating complex with α-actinin-4 and myosin IIb that pulls from the pointed-end toward the barbed-end of the actin filaments. This force generation results in the buckling of actin filaments and pulls on the plasma membrane in a manner that causes cells to extrude from a monolayer (24). One role for synaptopodin could be to function as a force-generating complex that pushes actin filaments into the protrusion structure and enables protrusion formation to be more efficient, but we anticipate this is unlikely to occur during infection. The barbed end of the filaments would be proximal to the bacteria where the incorporation of new actin monomers occurs. Thus, force generation by synaptopodin as described for the contractomere complex would generate a resistive force against protrusion formation by pulling the barbed end of actin back into the cell rather than pushing them out of the cell. Yet, we observe protrusions form faster with synaptopodin (Fig. 3E-G). An intriguing possibility is that synaptopodin, as part of the contractomere complex, is anchored at the plasma membrane and pulls the membrane along the actin filaments toward the bacteria. By pulling membrane into the protrusion, the contractomere complex would both create some slack in the membrane so as to enable the bacteria to more efficiently form the protrusion structure and would provide some mechanism for how the significant amount of plasma membrane is redirected into the protrusion. Although we have not seen previous descriptions of myosin IIb associating with bacteria during *S. flexneri* infection, previous reports do indicate roles for myosin IIa and myosin X (30, 50), and we hypothesize that myosin IIb could function in a similar manner by pulling the plasma membrane into the protrusion (30). A second possibility is that the presence of synaptopodin simply stabilizes the actin fibers, as described above, so that the loss of actin monomers is less efficient and the fibers remain longer in the protrusion for a greater period of time, thereby promoting the formation of the protrusion structure.

*Shigella* spp.*, Listeria* spp.*, Rickettsia* spp.*, Burkholderia* spp.*, Mycobacterium marinum,* and pox viruses such as vaccina virus are all examples of pathogens that require actin nucleation for their virulence. Recent studies show that mitochondria use similar mechanisms of motility by generating actin comet tails so as to enable organelle segregation during mitosis (51). Our findings show that synaptopodin is a critical regulator of actin polymerization and organization during *S. flexneri* and *L. monocytogenes* intercellular spread, and we anticipate that our findings represent a more general role of synaptopodin-dependent regulation of actin dynamics in cells and during infection.

## METHODS

### Bacterial Culture

*S. flexneri* strain 2457T is used in this study; all mutants are isogenic to it (8). *S. flexneri* strains were cultured in trypticase soy broth (TCS) at 37°C and at 250 rpm. Bacteria were transformed with pROEX-Aqua [Addgene; plasmid #42889, (52)] or pBR322-mCherry for live and fixed microscopy. *L. monocytogenes* strain 10403S is used in this study (53). *L. monocytogenes* strains were cultured in trypticase soy broth (TCS) at 37°C.

### Cell Culture

HeLa cells, MDCK cells (gift of Vivian Tang), and T84 cells were maintained at 37°C in 5% CO_2_. HeLa and MDCK cells were grown in Dulbecco’s modified Eagle’s Medium (DMEM) supplemented with 10% fetal bovine serum (FBS). T84 cells were grown in Advance DMEM supplemented with 10% FBS and 1% L-glutamine. The generation of cells producing control or synaptopodin specific shRNA were previously described for T84 cells (23) and MDCK cells (24). HeLa cells producing shRNA targeting synaptopodin were generated using retroviral transduction followed by selection with 5 µg/mL puromycin.

### Virus Preparation

3 million 293T cells were seeded in 100 mm dishes in DMEM supplemented with 10% FBS for 24 hours prior to transduction. Cells were transduced with psPAX2 (Addgene, 12260), pVSVg, and either pLKO.5 (SHC216) for control shRNA or a synaptopodin specific shRNA (TRCN0000218429, TRCN0000230911, TRCN0000230913, TRCN0000230912, TRCN0000218782), or pmbYFP (Gift of Hervé Agaisse). Plasmids were transfected into cells using calcium phosphate. The next day, the media was replaced with fresh DMEM supplemented with 10% FBS. After incubation for an additional 24 hours, the culture supernatant containing the viral particles was collected, passed through a 0.45 µm filter, and stored at −20°C.

### Viral Transduction

3 x 10^5^ HeLa cells in 6-well plates were treated with 2 mL of filtered 293 supernatant containing 4mM polybrene and centrifuged for 2 hours at 1000 *g* at 30°C. Cells were incubated at 37°C for 48 hours, and then treated with 5 µg/mL puromycin. Resistant cells were expanded. For cells additionally transduced with virus encoding the membrane anchored YFP, cells were sorted at the Flow Cytometry Shared Resource Laboratory in the Department of Immunology and Microbiology to enrich for YFP positive cells.

### Plaque Assay

8 x 10^5^ cells per well were seeded in 6-well plates. The next day, cells were infected with *S. flexneri* at a multiplicity of infection (MOI) of 0.2 for HeLa cells or a MOI of 20 for MDCK cells and T84 cells. Bacteria were centrifuged onto cells at 800 *g* for 10 minutes and incubated at 37°C in 5% CO_2_ for 90 min. Media was replaced with DMEM containing 10% FBS, 25 µg/mL gentamicin, and 0.5% agarose, and the cells were incubated for 48 hours. To visualize plaques, a second overlay was added with 0.7% agarose in DMEM with 10% FBS, 25 µg/mL gentamicin, 0.1% neutral red. Following incubation for at least 4 hours, the plates were imaged with an ImmunoSpot S6 Universal Visible/Fluorescent Analyzer (Cellular Technology Limited). Analysis of the plaque area was performed using a pipeline generated in ImageJ in which images were thresholded to remove background, pixel intensity was converted to a binary, and the images were segmented into objects; objects matching plaques on unmodified images were selected, and their area was quantified, as done previously (39). For experiments with butyrate, 5 mM butyrate was added 1 hour prior to infection and maintained throughout the infection.

### Spread Assay by Microscopy

Cells were seeded at 7 x 10^5^ cells per well in 6-well plates onto sterile coverslips. The next day, cells were infected with *L. monocytogenes* at a MOI of 15. The bacteria were centrifuged onto cells at room temperature at 800 *g* for 10 minutes. The co-culture was incubated at 37°C with 5% CO_2_ for 45 mins. Cells were washed and remaining extracellular bacteria were killed by DMEM supplemented with 10% FBS and 25 µg/mL gentamicin. The infection was incubated for an additional 24 hours and fixed with 3.7% paraformaldehyde. The cells were permeabilized with 1% triton X-100 in PBS and washed five times with PBS. Cells were stained with phalloidin conjugated to Alexa Flour 750 (Invitrogen A30105), washed with PBS, stained with Hoechst 33342 (Biotech No. 5117), washed twice with PBS, and mounted with Prolong Diamond Antifade Mountant (ThermoFisher, P36930). Images were taken at random at 20x magnification. Five fields of view were analyzed per condition per experiment and area of spread was measured with NIS elements.

### Quantification of Intracellular Bacteria

T84 cells were seeded at 8 X 10^5^ cells per well, and HeLa or MDCK cells were seeded at 4 X 10^5^ cells per well in a 6-well plate. The next day, cells were infected with *S. flexneri* at an MOI of 200, by centrifugation at at 800 *g* for 10 minutes. The co-culture was incubated at 37°C with 5% CO_2_ for 45 minutes. Excess bacteria were removed by three washes with HBSS. Extracellular bacteria were killed by DMEM supplemented with 10% FBS and 25 µg/mL gentamicin. The infection was incubated for an additional 1, 2, or 3 hours. The cells were washed three times with HBSS, and then the cells were lysed with 0.02% SDS in HBSS. The cell lysates with intact bacteria were serially diluted in TCS broth and plated onto TCS agar to determine the intracellular CFU.

### Western Blots

SDS-PAGE gels were transferred to nitrocellulose using a Turbo semi dry transfer apparatus (BioRad). For detection of antigen, 1:100 rabbit anti-synaptopodin (Abcam, ab224491) was incubated overnight at 4°C in 5% milk. The next day, blots were incubated at room temperature with 1:5,000 goat anti-rabbit (Jackson Immuno Research Labs, 111-035-144) for 2 hours or 1:10,000 anti-β-actin conjugated to HRP (Sigma A3854) for 1 hour. Western blots were developed, and the signal was acquired by a G:Box (Syngene). Band intensity was measured using ImageJ (NIH) with the background subtracted.

### Localization of Host Proteins

HeLa cells or MDCK cells were seeded at 8 X 10^5^ cells per well in a 6-well plate and cultured for 24 hours (HeLa) or 48 hours (MDCK). Cells were infected with designated strains of *S. flexneri* harboring pROEX-Aqua by centrifugation at 800 *g* for 10 minutes and incubated at 37°C in 5% CO_2_. At 45 minutes of infection, cells were washed three times with HBSS, and media was replaced with DMEM with 10% FBS, 25 µg/mL gentamicin, and 100 mM IPTG. After an additional 45 minutes, cells were fixed with 3.7% paraformaldehyde. The cells were permeabilized with 1% triton X-100 in PBS for 30 minutes at room temperature. The cells were washed five times with PBS and incubated in PBS with 10% FBS and 1:100 rabbit anti-synaptopodin (Abcam, ab224491), 1:1000 rabbit anti-N-WASP (Santa Cruz Biotechnology Inc, sc100964), 1:500 rabbit anti-α-actinin-4 (Abcam, ab108198), 1:50 rabbit anti-mDia1 (Abcam, AB129167), 1:500 mouse anti-myosin IIb (Santa Cruz Biotechnology Inc, sc-376942), or 1:50 goat anti-Arp 2 (Novus Biologicals, NB100-1037) overnight at 4°C. The cells were washed with PBS, incubated with 1:100 donkey anti-rabbit-Alexa Fluor 594 (ThermoFisher, A-21207), goat anti-rabbit-Alexa Fluor 750 (ThermoFisher, A21039), donkey anti-mouse-Alexa Fluor 594 (Invitrogen, A32744), or donkey anti-goat-Alexa Fluor 594 (ThermoFisher, A32758) for 2 hours at room temperature, and washed with PBS. Cells were stained with phalloidin-Alexa Flour 750 (Invitrogen A30105) or Alexa Fluor 647 (Invitrogen, A22287), washed with PBS, stained with Hoechst 33342 (Biotech No. 5117), washed twice with PBS, and mounted with Prolong Diamond Antifade Mountant (ThermoFisher, P36930).

### Live Imaging for Speed of Cytosolic Bacteria and of Protrusion Formation

HeLa cells expressing a membrane-anchored yellow fluorescent protein were seeded at 8 X 10^5^ cells per well on Ibidi 2-well chamber slides (Ibidi, 80286). The next day, cells were infected with designated strains of *S. flexneri* harboring the pROEX-Aqua plasmid at an MOI of 400 by centrifugation at 800 x*g* for 10 minutes. At 1 hour of infection, cells were washed with HBSS 10 times, and media was replaced with DMEM supplemented with 10% FBS, 25 µg/mL gentamicin, and 100 mM IPTG. A time-lapsed Z-stack of images was collected with a Nikon Eclipse Ti-2 inverted light microscope at 37°C and 5% CO_2_ at 10-second intervals for 10 minutes. Images were deconvolved using a Richardson-Lucy algorithm with 20 iterations and all slices within a stack were collapsed into a single image based upon the maximum intensity of pixels in each slice using NIS elements software. For time-lapsed series, in which some focal drift occurred in the XY position, FIJI plugin: Linear Stack Alignment with SIFT, was used to realign timepoints. NIS elements particle tracking software was used to track in-frame bacteria throughout the video to calculate protrusion velocities. Speed was determined for each bacterium by measuring the distance traveled over the duration of time bacterium was in focus.

### Live-cell Imaging Analysis for Protrusions Duration

HeLa cells expressing a membrane-anchored yellow fluorescent protein were seeded at 2 X 10^5^ cells per well on Ibidi 8-well chamber slides (Ibidi, 80826). The next day, media was replaced with DMEM and 10% FBS and cells were infected with designated strains of *S. flexneri* expressing mCherry at an MOI of 200. At 1 hour of infection, cells were washed with HBSS 10 times, and media was replaced with DMEM supplemented with 10% FBS. A time-lapsed Z-stack of images was taken with a Nikon Eclipse Ti-2 inverted light microscope at 37°C and 5% CO_2_ at 5-minute intervals for 2.5 hours to examine protrusion duration. Images were deconvolved using a Richardson-Lucy algorithm with 20 iterations and all slices within a stack were collapsed into a single image based upon the maximum intensity of pixels in each slice using NIS elements software. Protrusion duration was measured by quantifying the amount of time the bacteria were in each protrusion stage (54).

### Quantification of the length of actin tails formed by *L. monocytogenes*

HeLa cells were seeded at 8 X 10^5^ cells per well in a 6-well plate and cultured for 24 hours. The next day, cells were infected with *L. monocytogenes* harboring pGFP (55) at an MOI of 1,500. Bacteria were centrifuged onto cells at 800 *g* for 10 minutes and incubated at 37°C in 5% CO_2_. At 90 minutes of infection, cells were washed three times with HBSS, and media was replaced with DMEM with 10% FBS and 25 µg/mL gentamicin. After an additional 3.5 hours, cells were fixed with 3.7% paraformaldehyde for at least 20 minutes at room temperature. The cells were permeabilized with 1% triton X-100 in PBS for 30 minutes at room temperature. The cells were washed 5 times with PBS. Cells were stained with phalloidin conjugated to Alexa Flour 750 (Invitrogen A30105), washed with PBS, stained with Hoechst 33342 (Biotech No. 5117), washed twice with PBS, and mounted with Prolong Diamond Antifade Mountant (ThermoFisher, P36930).

### Microscopy

Fluorescent images were acquired on a Nikon Eclipse Ti-2 inverted light microscope equipped with an Orca Fusion BT cMOS camera (Hammamatsu), an IRIS 15 cMOS camera (Photometrics), Semrock Brightline filters, and an Oco stage top incubator chamber that enables regulation of temperature, CO_2_, and humidity. NIS elements software (Nikon) was used for image acquisition and analysis. Images were deconvolved using a Richardson-Lucy algorithm with 20 iterations. Microscopic images were pseudo-colored and assembled using Adobe photoshop or FIJI (NIH). Images were collected randomly across the coverslip using an automated imaging pipeline created in NIS elements or the samples were blinded prior to imaging. A minimum of five Z-stacks were collected per coverslip with 0.25 µm step size. To display representative images, all slices within a stack were collapsed into a single image based upon the maximum intensity of pixels in each slice using NIS elements software.

### Intensity Analysis

The signal intensity of random spots of synaptopodin, actin, mDia, Arp2, N-WASP, α-actinin-4 and myosin IIb around bacteria was measured using NIS elements. Three spots were analyzed at each bacterium, and 5-30 bacteria were measured per condition per experiment, and the average intensity was recorded per experiment.

### Statistical Analysis

Statistical difference between means was analyzed with Graph Pad Prism 9. Statistical difference of two means was determined either by Student’s t-test or a paired t-test. Unless indicated otherwise, the differences between the means of multiple groups were determined either by two-way ANOVA with Sidak *post hoc* test or by one-way ANOVA with either a Holm-Sidak’s *post hoc* test or a Dunnett’s *post hoc* test.

## Acknowledgements

We would like to thank Marcia Goldberg, Vivian Tang, Cammie Lesser, and Hervé Agaisse for reagents. We would like to thank Marcia Goldberg, Cristina Penaranda, Lilian Radoshevich, Marijke Keestra-Gounder and the Keestra-Gounder lab for helpful discussions. We would like to thank Tinalyn Kupfer and the CU | AMC ImmunoMicro Flow Cytometry Shared Resource, RRID:SCR_021321, for assistance with cell sorting. This work was funded by K22 AI137296, the GI and Liver Innate Immune Program at the University of Colorado, and the University of Colorado start-up funds to BCR. JER was funded by T32 AI052066.

## Author Contributions

J.M.V and B.C.R wrote the initial draft of the manuscript. J.M.V, D.J.H, T.B.H, T.N.H. S.P.C., L.L.L., and B.C.R designed experiments. J.M.V, J.D.T, B.C.R, D.J.H, T.B.H, T.N.H. L.Z.,and J.E.R preformed experiments. All authors discussed the results and commented on the manuscript.

## COMPETING FINANCIAL INTEREST

The authors declare no competing financial interest.

## Notes

### Competing Interest Statement

The authors have declared no competing interest.

### Summary of Updates

We provide additional data and clarify text.

