## Supplemental Material for "Synaptopodin is required for efficient intercellular spread by bacterial pathogens"

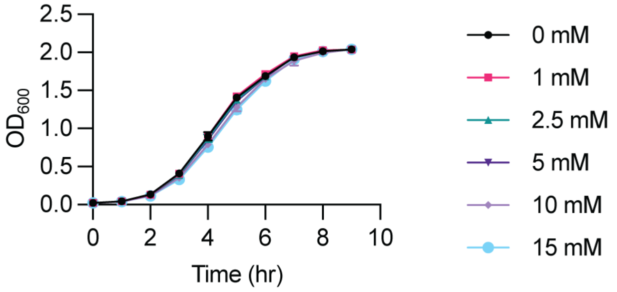


**Fig. S1: Butyrate does not alter *S. flexneri* replication.**

Bacteria were cultured in broth in the presence of various concentrations of butyrate. Data are mean ± SEM. Curves are not significantly different by non-linear regression.

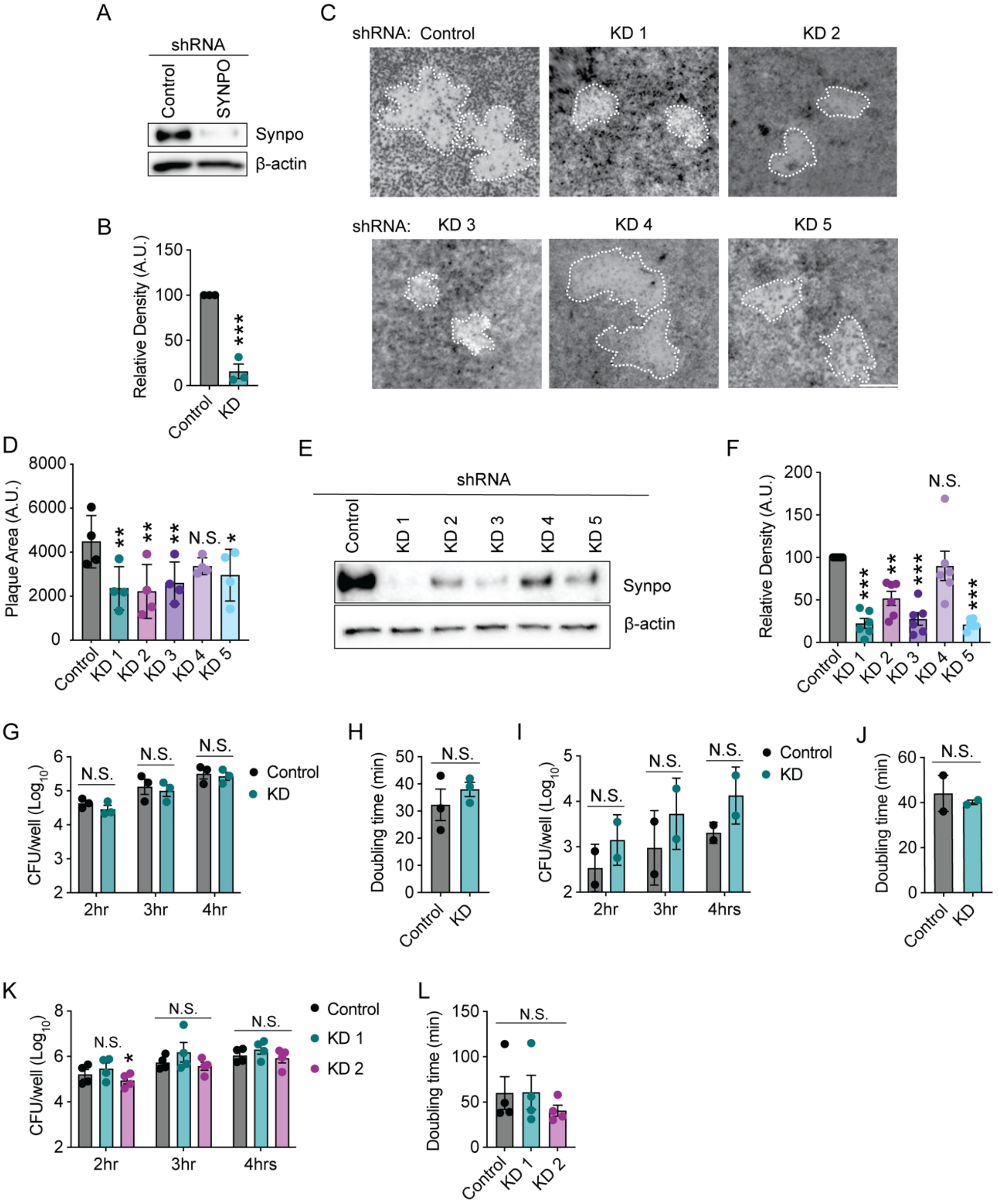


**Fig. S2. *S. flexneri* intracellular replication is not altered by loss of synaptopodin.**

(A) Western blot from lysates of MDCK cells producing a control shRNA or a shRNA specific to synaptopodin. (B) Quantification of relative of densitometry of synaptopodin from blots represented in A.*, $p<0.001$ by Student's t-test. (C) Representative images of HeLa cells infected with *S. flexneri* and stained with neutral red at 48 hours of infection. Control and KD1 images are reproduced in figure 2. Scale bar, 0.5 mm. Dotted lines indicate plaque boundary. (D) Quantification of plaques are from images represented in C. Quantification of Control, KD1, and KD2 are reproduced in figure 2. 15-60 plaques measured per condition per experiment. Dots are independent experiments. *, $p<0.05$; **, $p<0.01$; by one-way ANOVA with Dunnett's *post hoc* test. (E) Western blot from lysates of HeLa cells producing a control shRNA or a shRNA specific to synaptopodin. (F) Quantification of relative of densitometry of synaptopodin from blots represented in E. N.S., not significant. **, $p<0.01$,***, $p<0.001$ by one-way ANOVA with Dunnett's *post hoc* test. Dots are independent experiments. (G-L) Quantification of intracellular abundance of *S. flexneri* in T84 cells (G-H), in MDCK cells (I-J), and in HeLa cells (K-L). (G, I, and K) Quantification of colony forming units of bacteria recovered from cells. (H, J, and L) Quantification of the doubling time from 2 to 4 hours of infection. (B, D, F-L) N.S., not significant. *, $p<0.05$ by two-way ANOVA with Sidak *post hoc* test (G, I, and K), by one-way ANOVA with Dunnett's *post hoc* test (L) or by Student's t-test (B, H, and J). (B, D, F-L) Dot are independent experiments and data are mean $\pm$ SEM.

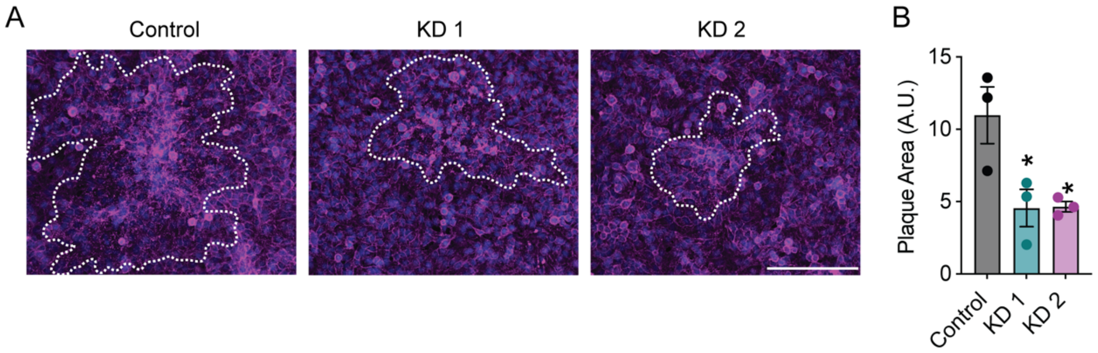


**Fig. S3. Synaptopodin is required for efficient intercellular spread of *L. monocytogenes.***

(A) Representative images of HeLa cells infected with *L. monocytogenes*. Magenta, actin; blue, DNA. Dotted lines indicate boundary of spread. Scale bar, 500 μM. (B) Quantification of plaque area from images represented in A. Five fields were analyzed per experiment. Dots are independent experiments. Data are mean ± SEM. *, $p<0.05$; by one-way ANOVA with Dunnett's *post hoc* test.

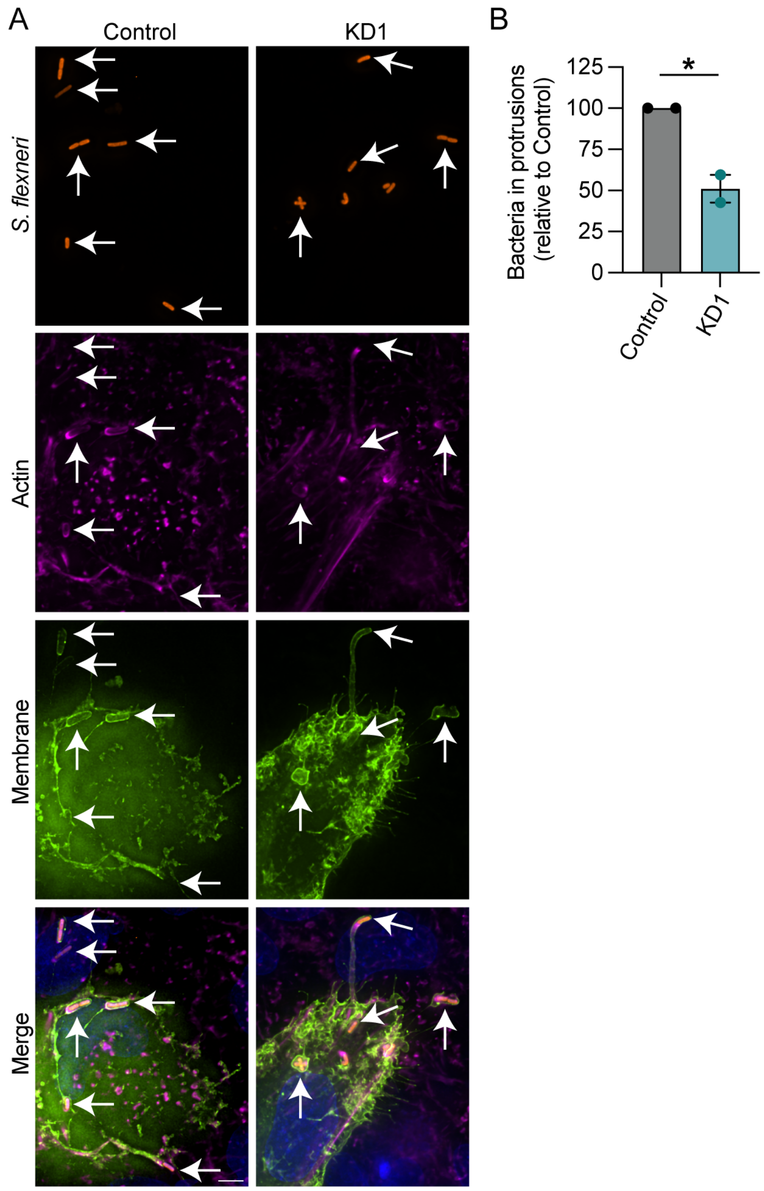


**Fig. S4. Protrusion formation is more efficient with synaptopodin.**

(A) Representative images of HeLa cells infected with *S. flexneri.* Orange, *S. flexneri*; magenta, actin; green, plasma membrane; blue, DNA. Arrows indicate bacteria in plasma membrane protrusions. Scale bar, 5 µM. (B) Quantification of the relative amount of bacteria in plasma membrane protrusions. *, $p<0.05$ by Student's t-test. Dots are individual experiments. Data are mean ± SEM.

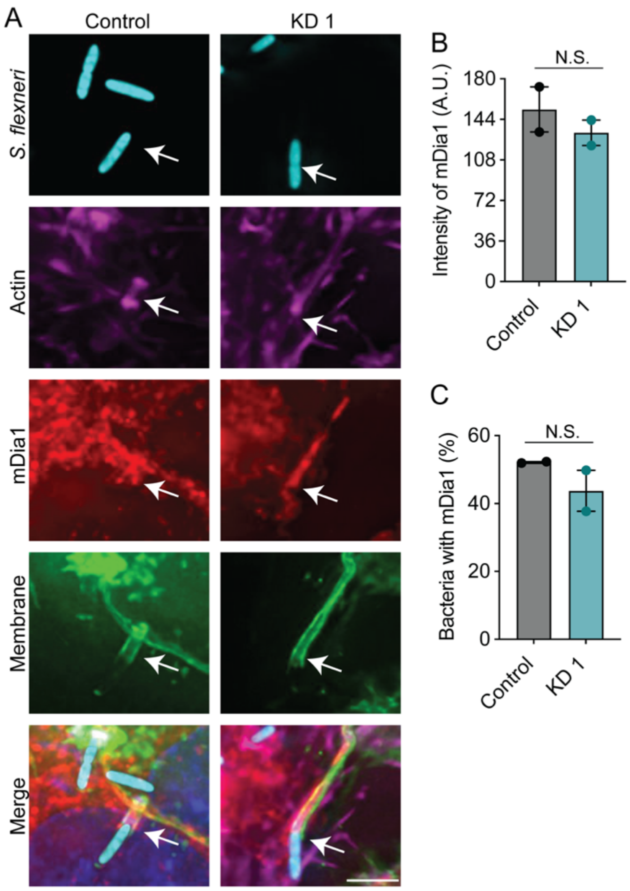


**Fig. S5. mDia1 is efficiently recruited to bacterial protrusions in the absence of synaptopodin.**

(A) Representative images of HeLa cells infected with *S. flexneri.* Cyan, *S. flexneri;* magenta, actin; red, mDia1; green, plasma membrane; blue, DNA. Arrows, bacteria in a plasma membrane protrusion. Scale bar, 5 μM. (B) Quantification of the mDia1 intensity in the protrusions structure. (C) Quantification of percentage of bacteria with mDia1 recruitment. (B and C) N.S., not-significant by Student's t-test. Dots are independent experiments. Data are mean ± SEM.

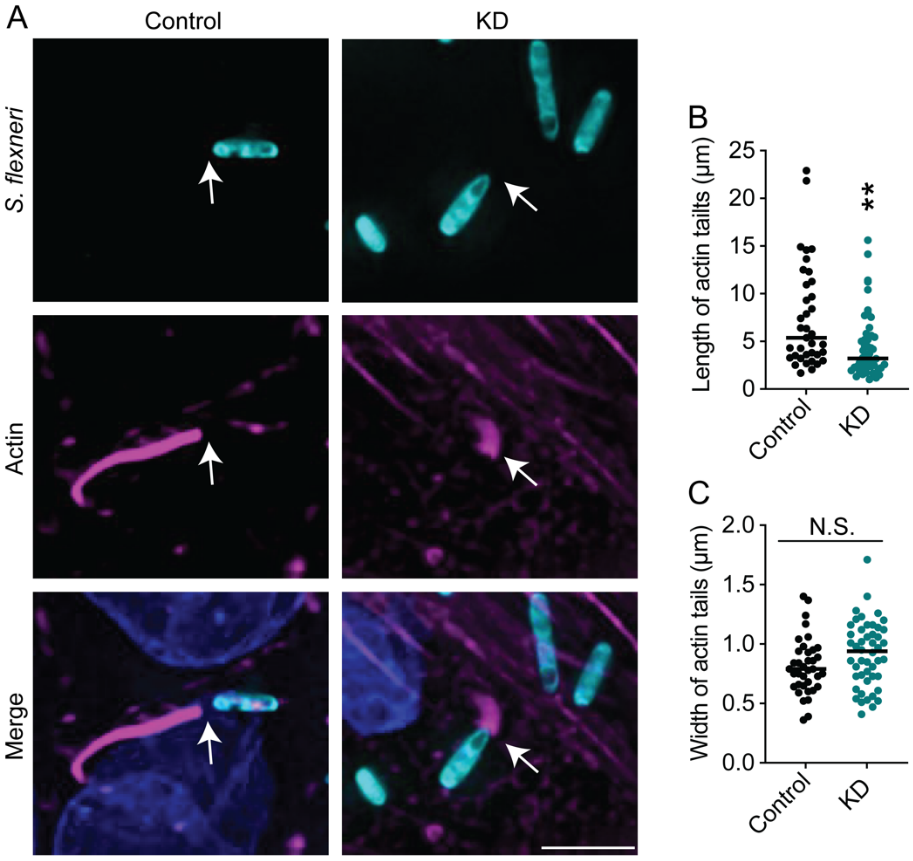


**Fig. S6. Synaptopodin regulates actin tail formation by *S. flexneri* in MDCK cells.**

Infection of MDCK cells with *S. flexneri* for 8 hours. (A) Representative images. Cyan, *S. flexneri;* magenta, actin; blue, DNA. Arrows, bacteria with actin tails. Scale bar, 5 µM. (B) Quantification of the length of actin tails. (C) Quantification of the width of actin tails proximal to the bacterial pole. (B and C) 5-10 fields were analyzed per experiment. Dots are individual bacteria pooled together from three experiments. Data are mean ± SEM. N.S., not significant; **, p<0.01; by Student's t-test.

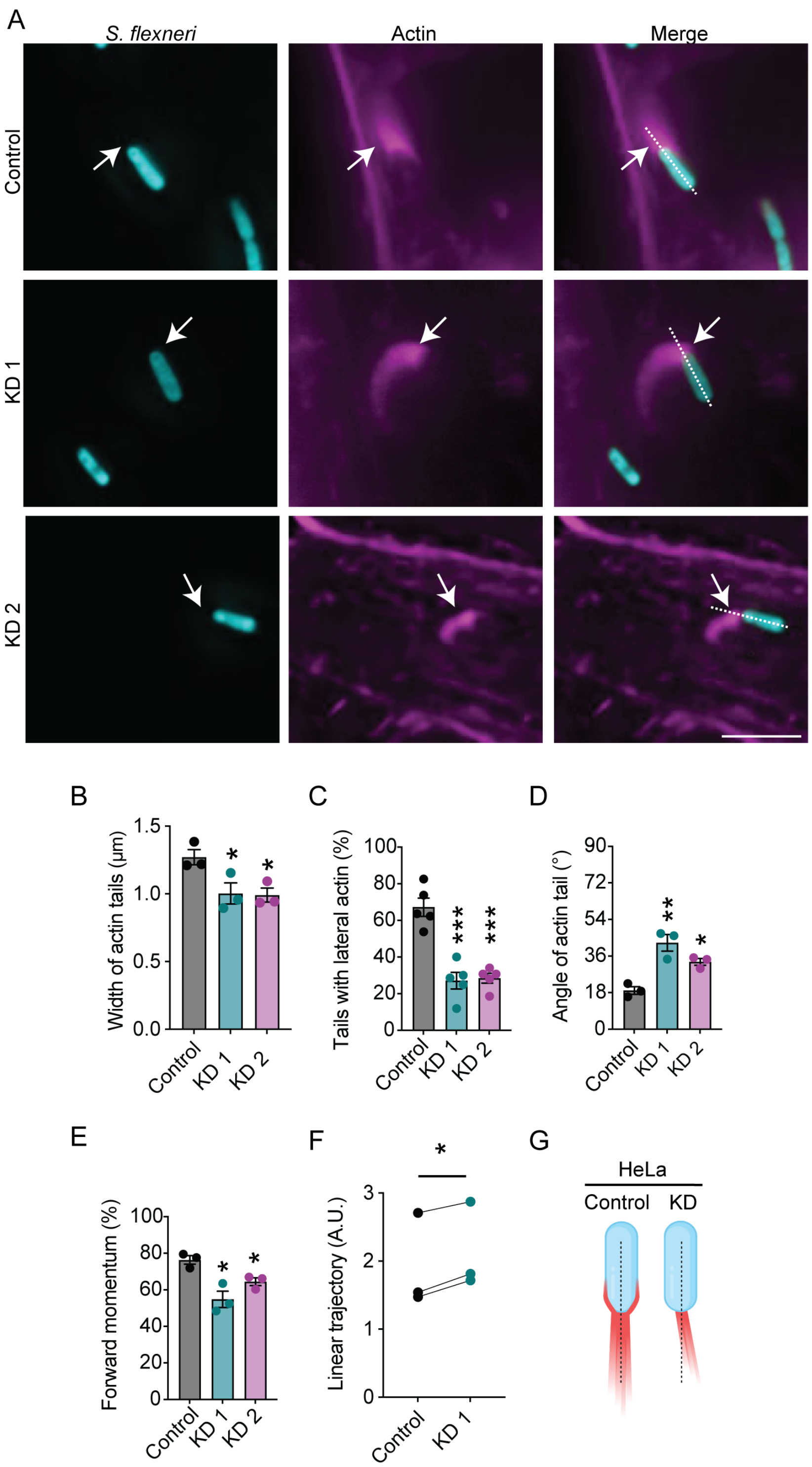


64

65 **Fig. S7: Additional actin tail phenotypes observed in HeLa cells with synaptopodin**

66 **knockdown.**

Infection of HeLa cells infected with *S. flexneri* for 90 min. A) Representative images. Cyan, *S. flexneri;* magenta, actin; blue, DNA. Arrows, bacteria with actin tails. Scale bar, 5 μM. (B) Quantification of the width of actin tails proximal to the bacterial pole. (C) Quantification of the percent of bacteria with an actin tail that also display lateral actin from images represented in A. (D) Quantification of the angle (0°-90°) at which the actin tail is anchored to the bacterial pole with respect to the longitudinal midline of the bacteria from images represented in A. (E) Mathematical estimation of the forward momentum as a result of the angle of the tail from images of bacteria represented in A. 100% forward moment corresponds to a tail angle of 0°. (F) Quantification of the linearity of the path bacteria traveled in the cell from videos represented in Fig. 3D. (G) Schematic of actin tail for bacteria in control or synaptopodin knockdown HeLa cells. Created with biorender.com. (B-E) 5-10 fields were analyzed per experiment. Dots are independent experiments. Data are mean ± SEM. N.S., not significant; *, $p<0.05$; **, $p<0.1$; ***, $p<0.001$ by one-way ANOVA with Dunnett's *post hoc* test. (F) N.S., not significant; *, $p<0.05$ by paired t-test. (B-F) Dots are means of individual experiments.

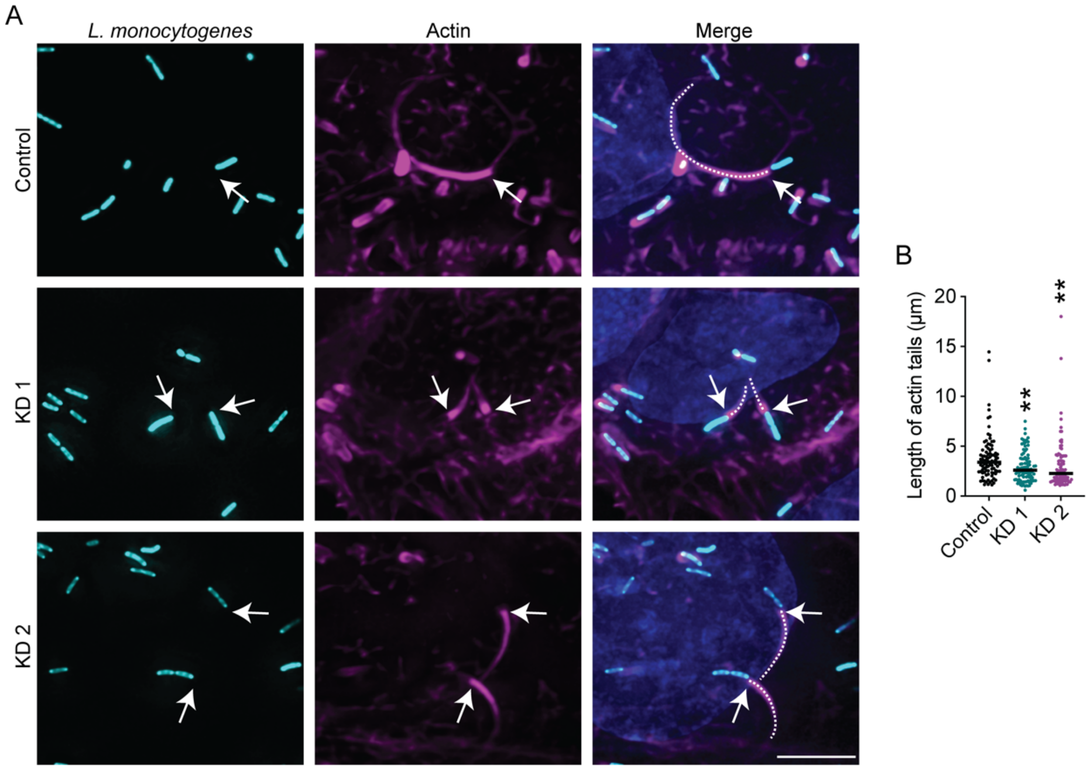


**Fig. S8: Actin tails formed by *Listeria monocytogenes* are longer with synaptopodin.**

HeLa cells infected with *L. monocytogenes* for 5 hours. (A) Representative images. Cyan, *L. monocytogenes;* magenta, actin; blue, DNA. Arrows, bacteria with actin tails. Dotted line represents actin tail length. Scale bar, 5 µM. (B) Quantification of the length of the actin tails. Dots are individual bacteria pooled together from three experiments. **, p>0.01; by Kruskal-Wallis test with Dunn's *post hoc* test.

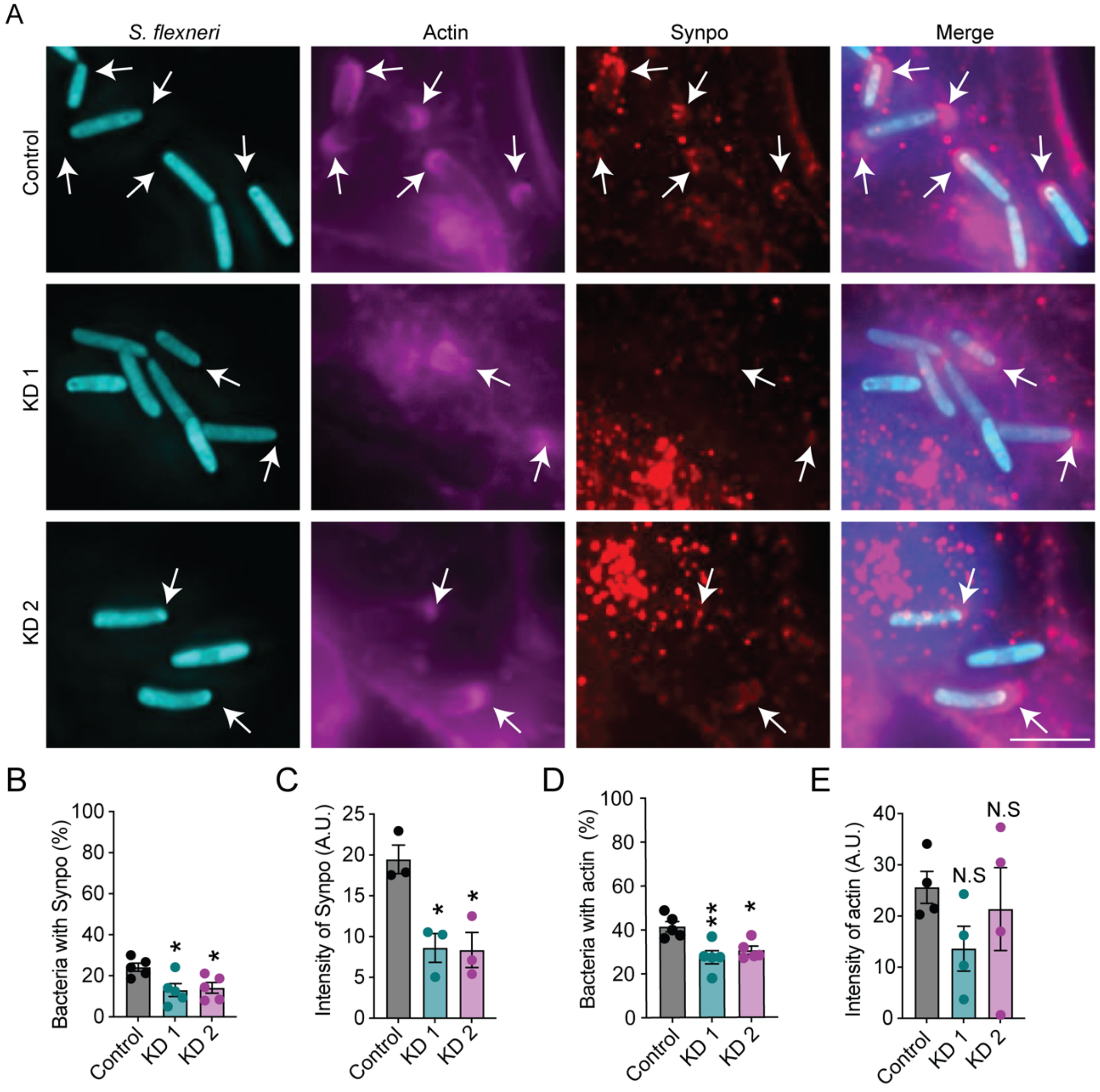


**Fig. S9. Synaptopodin is recruited to bacteria with actin.**

HeLa cells infected with *S. flexneri* for 90 minutes. (A) Representative images of HeLa cells infected with *S. flexneri.* Cyan, *S. flexneri;* magenta, actin; red, synaptopodin; blue, DNA. Arrows, bacteria with actin and synaptopodin. Scale bar, 5 μM. (B-C) Dots are independent experiments. (B) Quantification of percentage of bacteria that recruit synaptopodin. (C) Quantification of intensity of synaptopodin localized around bacteria. (D) Quantification of percentage of bacteria with actin. (E) Quantification of intensity of actin localized around bacteria. (B-E) 5-10 fields were analyzed per condition per experiment. Dots are independent experiments. Data are mean ± SEM. N.S., not significant; $p>0.05$; *, $p<0.05$; **, $p<0.01$ by one-way ANOVA with Dunnett's *post hoc* test.

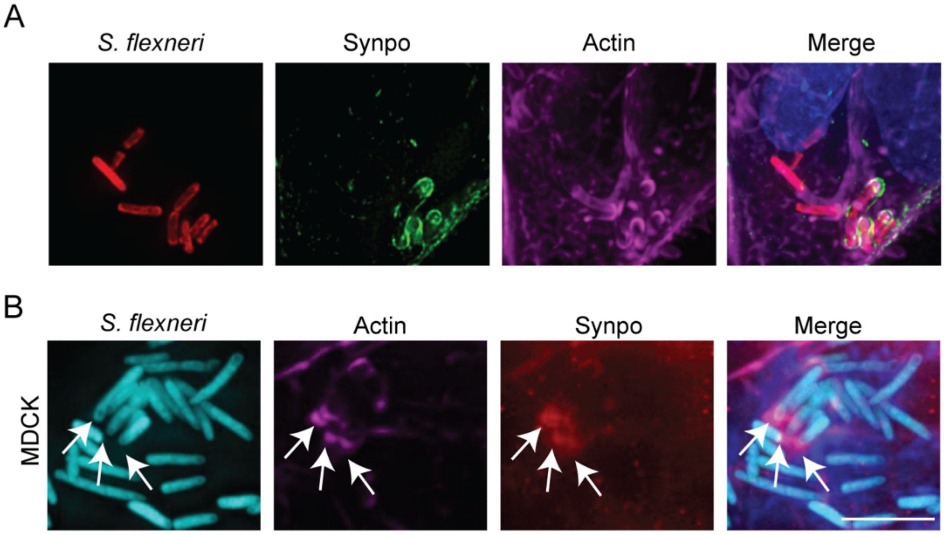


**Fig. S10. Synaptopodin is recruited around *S. flexneri* in MDCK cells.**

(A) Representative images of HeLa cells producing synaptopodin-venus and infected with *S. flexneri* for 90 minutes. Green, synaptopodin; magenta, actin; red, *S. flexneri;* blue, DNA. (B) Representative images of MDCK cells infected with *S. flexneri.* Cyan, *S. flexneri;* magenta, actin; red, synaptopodin; blue; DNA.

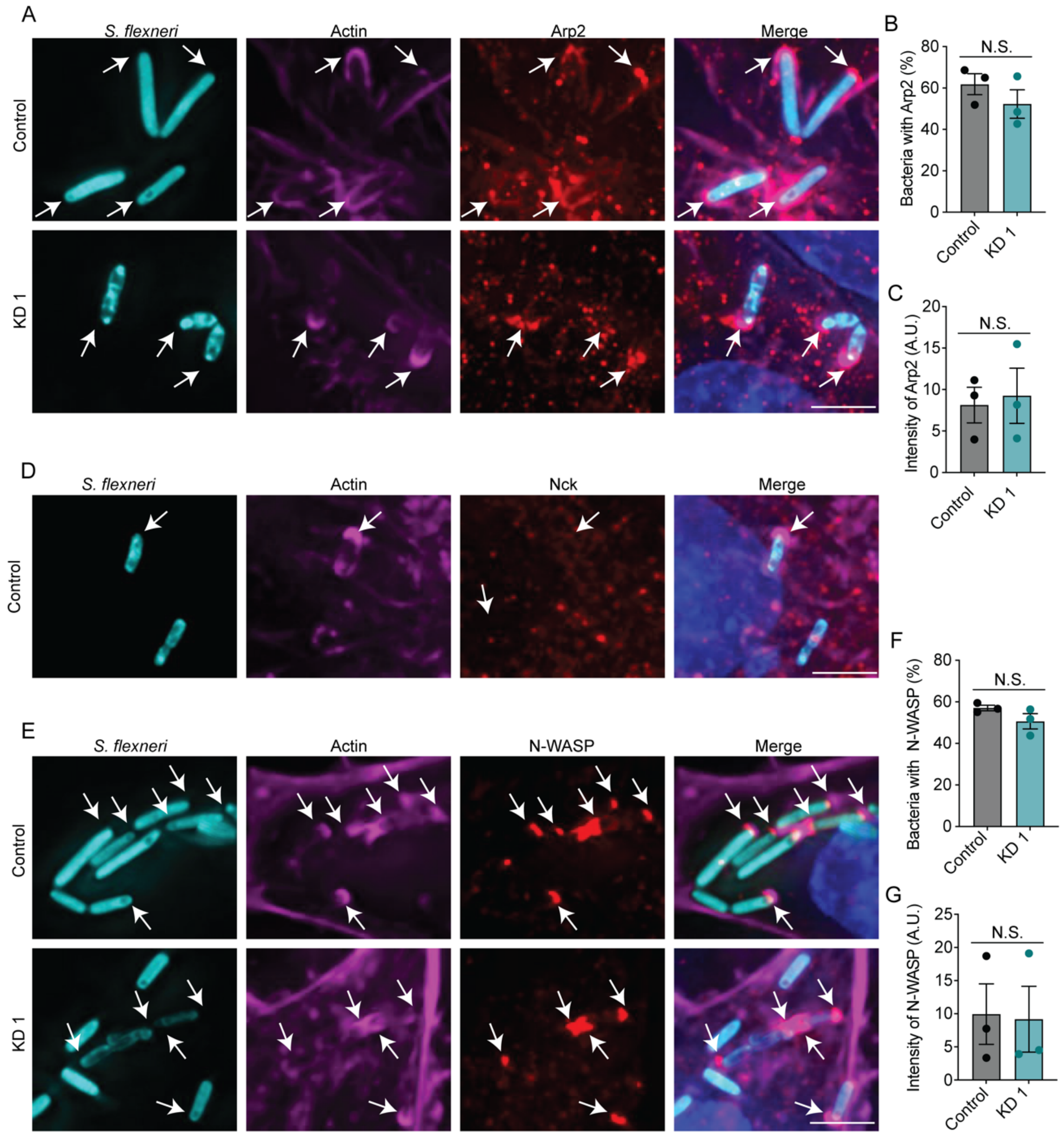


**Fig. S11. Nck1, Arp2, and N-WASP are recruited to bacteria in the relative absence of synaptopodin.**

HeLa cells infected with *S. flexneri* for 90 min and tested for the recruitment of Arp2 (A-C), Nck-1 (D), or N-WASP (E-G) around bacteria. (A) Representative images depicting Arp2 around bacteria. Cyan, *S. flexneri;* magenta, actin; red, Arp2; blue, DNA. Arrows, bacteria with actin and Arp2. Scale bar, 5 μM. (B) Quantification of the percentage of bacteria with Arp2. (C)

Quantification of the intensity of Arp2 around bacteria. (D) Representative images depicting lack of NCK-1 recruitment around bacteria. Cyan, *S. flexneri;* magenta, actin; red, NCK1; blue, DNA. Arrows, bacteria with actin. Scale bar, 5 μM. (E) Representative images depicting N-WASP recruitment around bacteria. Cyan, *S. flexneri;* magenta, actin; red, N-WASP; blue, DNA. Arrows, bacteria with actin and N-WASP. Scale bar, 5 μM. (F) Quantification of the percentage of bacteria with N-WASP. (G) Quantification of the intensity of N-WASP around bacteria. (B, C, F, and G) N.S., not significant by paired t-test. Dots are independent experiments. Data are ± SEM.

**Movie S1: Time lapse videos of bacteria forming plasma membrane protrusions in control cells.**

*S. flexneri* infection of control HeLa cells. Cyan, S. *flexneri;* green, plasma membrane. Images taken at 10-second intervals for 10 minutes.

**Movie S2: Time lapse videos of bacteria forming plasma membrane protrusions in Synpo KD cells.**

*S. flexneri* infection of synaptopodin knockdown HeLa cells. Cyan, S. *flexneri;* green, plasma membrane. (Images taken at 10-second intervals for 10 minutes.

**Movie S3: Time lapse videos of intercellular motility of bacteria in control cells.**

*S. flexneri* infection of control HeLa cells. Cyan, S. *flexneri;* green, plasma membrane. Images taken at 10-second intervals for 10 minutes.

**Movie S4: Time lapse videos of intercellular motility of bacteria in Synpo KD cells.**

*S. flexneri* infection of synaptopodin knockdown HeLa cells. Cyan, S. *flexneri;* green, plasma membrane. Images taken at 10-second intervals for 10 minutes.
